# Pancreatic islet α cell function and proliferation requires the arginine transporter SLC7A2

**DOI:** 10.1101/2023.08.10.552656

**Authors:** Erick Spears, Jade E. Stanley, Matthew Shou, Linlin Yin, Xuan Li, Chunhua Dai, Amber Bradley, Katelyn Sellick, Greg Poffenberger, Katie C. Coate, Shristi Shrestha, Regina Jenkins, Kyle W. Sloop, Keith T. Wilson, Alan D. Attie, Mark P. Keller, Wenbiao Chen, Alvin C. Powers, E. Danielle Dean

**Affiliations:** Division of Diabetes, Endocrinology, and Metabolism, Department of Medicine, Vanderbilt University Medical Center, Nashville, TN; Department of Biology, Belmont University, Nashville, TN; Department of Molecular Physiology & Biophysics, Vanderbilt University, Nashville, TN; Department of Biochemistry, University of Wisconsin, Madison, WI; Diabetes and Complications, Lilly Research Laboratories, Eli Lilly and Company, Indianapolis, IN; Division of Gastroenterology, Hepatology and Nutrition, Department of Medicine, Vanderbilt University Medical Center, Nashville, TN; Department of Pathology, Microbiology and Immunology, Vanderbilt University Medical Center, Nashville, TN; Center for Mucosal Inflammation and Cancer, Vanderbilt University Medical Center, Nashville, TN; Program in Cancer Biology, Vanderbilt University School of Medicine, Nashville, TN; Veterans Affairs Tennessee Valley Healthcare System, Nashville, TN

## Abstract

Interrupting glucagon signaling decreases gluconeogenesis and the fractional extraction of amino acids by liver from blood resulting in lower glycemia. The resulting hyperaminoacidemia stimulates α cell proliferation and glucagon secretion via a liver-α cell axis. We hypothesized that α cells detect and respond to circulating amino acids levels via a unique amino acid transporter repertoire. We found that *Slc7a2ISLC7A2* is the most highly expressed cationic amino acid transporter in α cells with its expression being three-fold greater in α than β cells in both mouse and human. Employing cell culture, zebrafish, and knockout mouse models, we found that the cationic amino acid arginine and SLC7A2 are required for α cell proliferation in response to interrupted glucagon signaling*. Ex vivo* and *in vivo* assessment of islet function in *Slc7a2^−/−^* mice showed decreased arginine-stimulated glucagon and insulin secretion. We found that arginine activation of mTOR signaling and induction of the glutamine transporter SLC38A5 was dependent on SLC7A2, showing that both’s role in α cell proliferation is dependent on arginine transport and SLC7A2. Finally, we identified single nucleotide polymorphisms in *SLC7A2* associated with HbA1c. Together, these data indicate a central role for SLC7A2 in amino acid-stimulated α cell proliferation and islet hormone secretion.

## Introduction

Pancreatic α cells secrete glucagon which stimulates gluconeogenesis and glycogenolysis in the liver when blood glucose concentrations are low. When glucose levels rise, β cells secrete insulin which counteracts these effects. The counterregulatory relationship between these two hormones is essential to maintenance of glucose homeostasis. The development of type 2 diabetes (T2D) results from reduced insulin secretion from pancreatic islets, and diminished insulin action in peripheral tissues. However, hyperglucagonemia in T2D, due to the inability to suppress glucagon secretion, also promotes T2D-associated hyperglycemia, highlighting a central role for α cell dysregulation in addition to insulin insufficiency (1). Interrupting glucagon action lowers blood glucose under non-diabetic conditions and in rodent models and humans with diabetes (2–5). Therefore, glucagon receptor antagonism is under investigation as a therapeutic approach to treat both type 1 diabetes (T1D) and T2D (6–8). Studies of glucagon antagonism led to the discovery of an interaction between the α cells in the pancreatic islet and hepatocytes that has been termed the liver-α cell axis. Interrupted glucagon signaling decreases hepatic amino acid extraction from the circulation and catabolism resulting in hyperaminoacidemia. This in turn feeds back to α cells, stimulating them to proliferate and secrete more glucagon. This response to interrupting glucagon signaling by any approach, is conserved in zebrafish, mouse, and human islets (9–14).

Most recently, studies have identified glutamine as being important for this amino acid-stimulated α cell proliferation and involves the mTOR-dependent upregulation of expression of the glutamine transporter, *Slc38a5* (9, 10). These studies also indicated roles for other amino acids in the liver-α cell axis as elevated levels of glutamine were essential, but not sufficient, suggesting a role for a combination of other amino acids like those found in serum of mice with interrupted glucagon signaling (10). Furthermore, global loss of *Slc38a5* expression reduced, but did not completely prevent, α cell proliferation following glucagon signaling interruption (9).

Arginine has profound physiological effects on islet cells as a potent secretagogue for both insulin and glucagon (15, 16). The most common arginine transporters belong to the Slc7a subfamily of y^+^-type cationic amino acid transporters historically known as CAT proteins (SLC7A1-4 and SLC7A14 (17)). Of all these subfamily members, *SLC7A2* is the most highly expressed in human pancreas and liver (gtexportal.org). In mice, *Slc7a2* expression was decreased in the liver in response to interrupted glucagon signaling and that this was accompanied by an increase in serum arginine (9, 10). Though SLC7A2 and arginine transport have been studied in other tissues and cell types, including macrophages, astrocytes, lung, and intestine (18–21), its role in islet cell physiology has not been assessed.

These observations led us to hypothesize that arginine plays a central role in the liver-α cell axis and that SLC7A2 is the primary arginine transporter in α cells. Here, we show that *Slc7a2* is highly expressed in α and β cells from humans, mice and zebrafish and that arginine is required for amino acid-stimulated α cell proliferation. Using genetic loss-of-function models in zebrafish and mice, we demonstrate that SLC7A2 is required for amino acid-stimulated α cell proliferation following interrupted glucagon signaling and that it plays a critical role in arginine-stimulated insulin and glucagon secretion from pancreatic β or α cells, respectively. Finally, we demonstrated that *Slc7a2* expression is required for the upregulation of *Slc38a5* expression following interrupted glucagon signaling. Together these studies reveal a conserved role for the arginine transporter, SLC7A2, in amino acid-regulated islet cell biology.

## Results

### Arginine is necessary for amino acid-stimulated α cell proliferation *in vitro*

Previous studies demonstrated the importance of hyperaminoacidemia for α cell proliferation and specifically a role for glutamine and its major transporter, SLC38A5 in mice and zebrafish (9, 10). Since *Slc38a5* expression differs between mouse and human α cells and its ablation only partially suppressed α cell proliferation (9, 10), we evaluated whether other amino acids are individually required for amino acid-stimulated α cell proliferation, as is glutamine. As described previously, culturing isolated mouse islets in high amino acid medium increased α cell proliferation (Fig 1A) (10). This high amino acid medium (All AA +) mimicked levels in mice with interrupted glucagon signaling while low amino acid medium (All AA -) was similar to levels in wildtype mouse serum (Supplemental Table 1). To identify other critical amino acids, we adopted a two-step approach to minimize the number of islets in the screen. As a first step, we reduced the levels of groups of 4 amino acids in the high amino acid medium (Supplemental Table 1). Only islets cultured in medium with low arginine, histidine, glycine, and asparagine (RHGN^low^) had decreased α cell proliferation (Fig 1A). To determine if a low level of a single amino acid in the RHGN^low^ medium was responsible, we then individually replaced each of these four amino acids in the RHGN^low^ medium at the high concentration and found that only arginine (R) stimulated amino acid-stimulated α cell proliferation (Fig 1B). We also assessed the importance of arginine for proliferation of a cultured α cell line, αTC1-6 cells. Culturing these cells in their normal DMEM-based medium lacking arginine inhibited proliferation (Fig 1C, Supplemental Table 2). These results indicate that arginine in addition to glutamine, is required for α cell proliferation (10).

**Figure 1.**
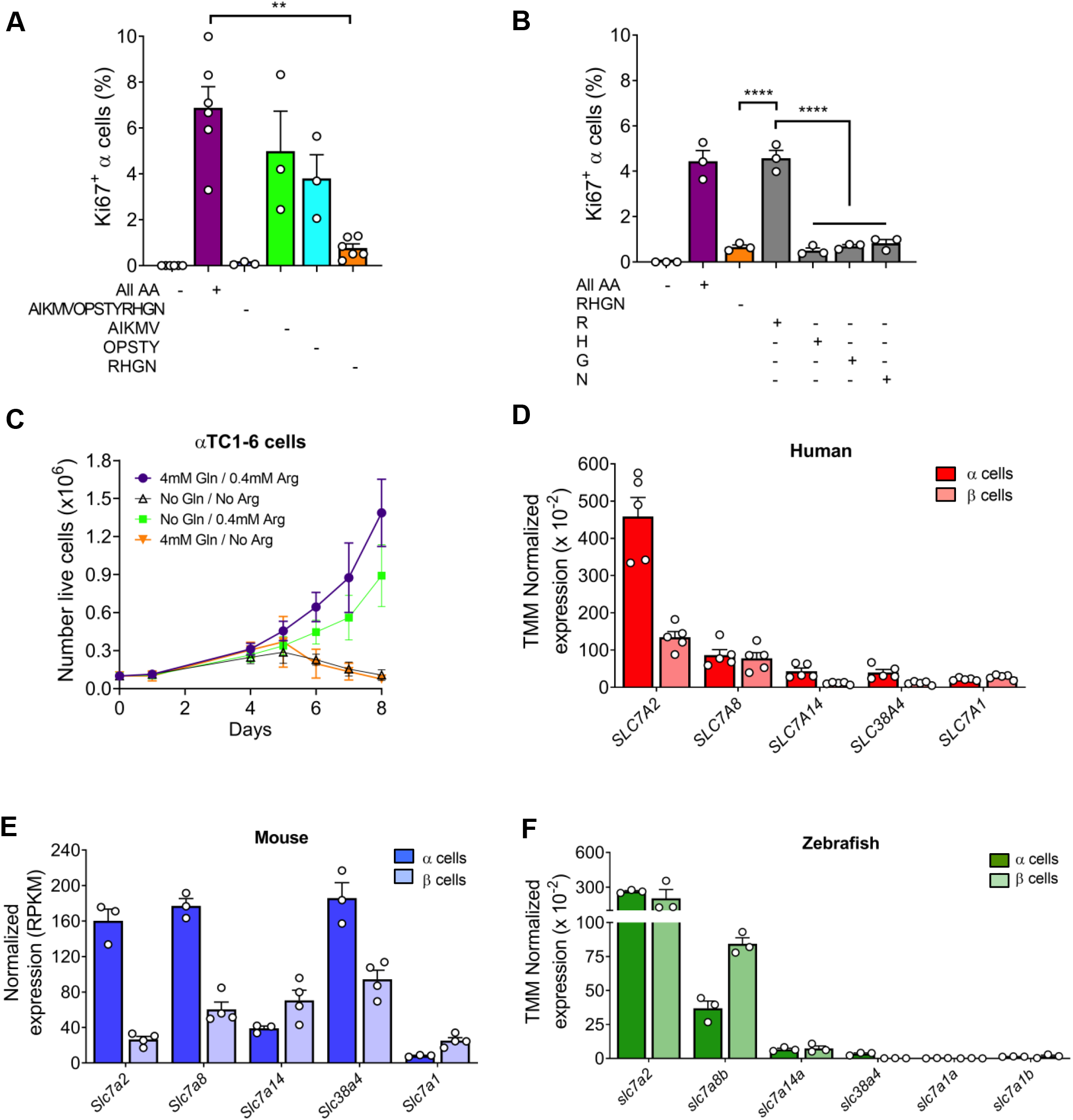
Arginine stimulates α cell proliferation in cultured mouse islets and cationic amino acid transporter, *Slc7a2*, expression is conserved in human, mouse, and zebrafish α cells. (**A**) *Ex vivo* α cell proliferation analysis in low (All AA -) and high (All AA +) amino acid containing medium, medium with low concentrations of 14 of the 20 total amino acids typically added to culture medium (AIKMVOPSTYRHGN -), otherwise high AA medium with Alanine (A), Isoleucine (I), Lysine (K), Methionine (M), and Valine (V) at low concentrations (AIKMV -), otherwise High AA medium with Hydroxyproline (O), Proline (P), Serine (S), Threonine (T), and Tryptophan (Y) at low concentrations (OPSTY -) and otherwise high AA medium with Arginine (R), Histidine (H), Glycine (G) and Asparagine (N) at low concentrations (RHGN -). (**B**) *Ex vivo* α cell proliferation analysis in low (-) and high (+) amino acid containing medium, in otherwise high AA medium with Arginine (R), Histidine (H), Glycine (G) and Asparagine (N) at low concentrations (orange bar, RHGN -), and with each of the low concentratin amino acids in low RHGN medium restored to high levels (gray bars, R +, H +, G + and N +). α cell proliferation was determined by percent Ki67+/Gcg+ cells per total Gcg+ cells in isolated islets treated cultured in medium containing different variations of amino acid concentrations (n=3 per group). ΑA = all amino acids, R = arginine, H = histidine, G = Glycine, N = asparagine, + = high AA concentrations (equivalent to that in serum of *Gcgr*^−/−^ mice), - = low AA concentration (equivalent to that in serum of *Gcgr*^+/+^). (**C**) Cell growth over time of αTC1-6 cultured cells in control DMEM or in DMEM lacking glutamine (Gln), arginine (Arg) or both. (**D-F**) Comparison of cationic amino acid transporter expression in α and β cells from published human (n=5; (22, 23)), mouse (n=3-4; (24) and Zebrafish (n=3; (25)) RNAseq datasets. Human and Zebrafish expression data were normalized with TMM method, mouse data were normalized to RPKM. Expression of other cationic amino acid transporter not shown were below the limits of detection in islet cells. Significance was designated: *p<0.05, **p<0.005, ***p<0.0005 and ****p<0.0001. Created with BioRender.com.

### *Slc7a2* is highly expressed in pancreatic α and β cells

Because interruption of glucagon signaling leads to increased serum amino acids and increased α cell proliferation (9, 10), we evaluated the expression of cationic amino acid transporters in pancreatic α and β cells by mining published transcriptomics datasets. In human RNAseq datasets (22, 23), *SLC7A2* is the most highly expressed of all SLC (<u>S</u>o<u>L</u>ute <u>C</u>arrier) superfamily genes in α cells and its expression is three-fold higher in α cells than in β cells (Fig 1D, Supplemental Table 3). From mouse RNA-seq data (24), *Slc7a2* is one of the most highly expressed amino acid transporters with a six-fold greater expression in α cells when compared to β cells (Fig 1E, Supplemental Table 3). In zebrafish (25), *slc7a2* is the third most highly expressed amino acid transporter (Supplemental Table 3) but is the most highly expressed cationic amino acid transporter (Fig 1F). The greater level of *Slc7a2* expression in islet cells of humans, mice, and zebrafish points toward its evolutionarily conserved importance in the endocrine pancreas and α cells.

### *Slc7a2^−/−^* mice have decreased arginine-stimulated glucagon and insulin secretion

To understand the role of arginine transport in islet function and α cell proliferation, we examined the impact of loss of *Slc7a2* on glucose homeostasis and islet function using a global *Slc7a2* knockout mouse line (*Slc7a2^−/−^*) (18-20, 26). We found that *Slc7a2^−/−^* mice had normal glucose tolerance in response to intraperitoneal glucose injection (Supplemental Fig. 1A and B). To assess stimulated glucagon secretion, *Slc7a2^−/−^* mice were given an intraperitoneal glucose/arginine bolus and serum glucagon concentrations measured. Stimulated blood glucose levels were higher in *Slc7a2^−/−^* than in *Slc7a2^+/+^* mice (Fig. 2A) supporting a role for SLC7A2 in glycemic regulation. Serum glucagon levels increased similarly in *Slc7a2^+/+^* and *Slc7a2^+/−^* after stimulation with glucose and arginine. However, stimulated glucagon levels in *Slc7a2^−/−^* mice were 60% lower than their *Slc7a2^+/+^* and *Slc7a2^+/−^*littermates and not different from fasted glucagon levels in the same *Slc7a2^−/−^*mice (Fig. 2B). Similar to glucose/arginine bolus, intraperitoneal administration of arginine alone did not increase serum glucagon in *Slc7a2^−/−^* animals as stimulated values were 95% lower than *Slc7a2^+/+^* mice and not significantly different from fasted glucagon values in the same mice (Fig. 2E), indicating a central role for SLC7A2 in arginine-stimulated glucagon secretion.

**Figure 2.**
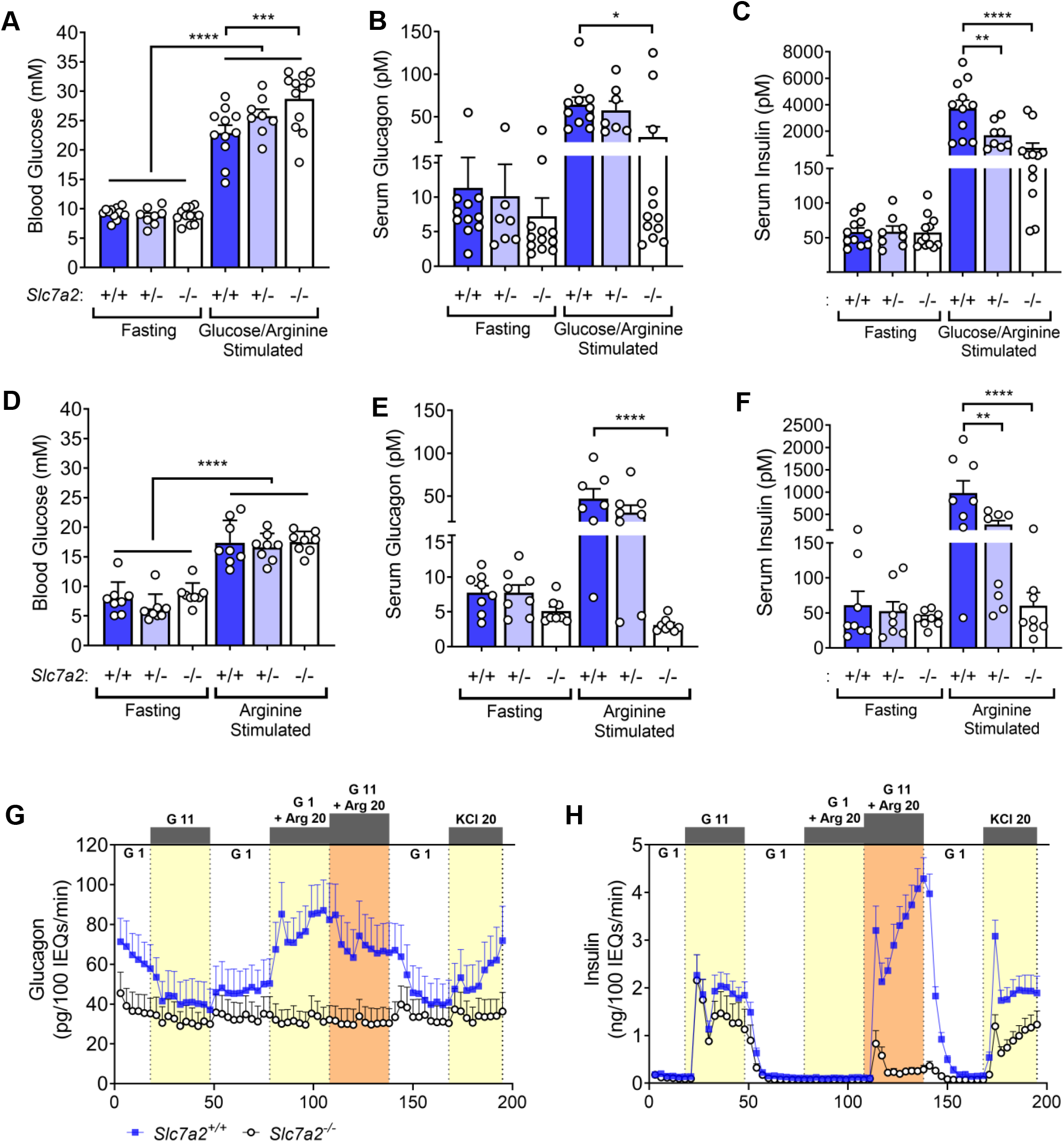
*Slc7a2^−/−^* mice have an impaired arginine-stimulated glucagon and insulin response *in vivo*. (**A**) Blood glucose, **(B)** serum glucagon, and **(C)** serum insulin for *Slc7a2^+/+^*(n=11), *Slc7a2^+/−^* (n=8) and *Slc7a2^−/−^* (n=12) mice fasted for 6 hours, injected with *glucose/arginine* bolus and sampled 15 minutes post injection. **(D)** Blood glucose, **(E)** serum glucagon, and **(F)** serum insulin for *Slc7a2^+/+^*, *Slc7a2^+/−^*and *Slc7a2^−/−^* mice fasted for 6 hours, injected with *arginine* bolus and sampled 15 minutes post injection (n=8 each genotype). (**G**) Glucagon and **(H)** insulin secretion as assessed by perifusion of islets isolated from *Slc7a2^+/+^* and *Slc7a2^−/−^* mice (n=4 each genotype). G 1 = 1 mM glucose, G 11 = 11 mM glucose, Arg 20 = 20 mM arginine, KCl 20 = 20 mM KCl. Total glucagon and insulin content of perifused islets are shown in Supplementary Figure 1 G, H.

Interestingly, glucose/arginine-stimulated serum insulin levels were similarly decreased in *Slc7a2^−/−^* animals, but there was also a decrease in glucose/arginine-stimulated serum insulin in *Slc7a2^+/−^* animals (Fig. 2C). Arginine did not increase serum insulin levels in *Slc7a2^−/−^* mice and arginine-stimulated insulin levels were also lower in *Slc7a2^+/−^* mice (Fig. 2F). These data indicate that arginine transport via SLC7A2 is necessary for arginine-stimulated glucagon and insulin secretion. Of note, the decrease in arginine-stimulated insulin secretion in *Slc7a2^+/−^* mice indicate that arginine transport may be the rate limiting step in stimulated insulin secretion.

### *Slc7a2^−/−^* mouse islets have an islet-intrinsic impaired response to arginine resulting in defective glucagon secretion

Because we used a global *Slc7a2* knockout mouse, we assessed whether the observed defects in arginine-stimulated secretion *in vivo* were due to intrinsic islet SLC7A2 loss (islet autonomous) or mediated by extra-islet signals by perifusing isolated islets from *Slc7a2^−/−^*mice, and *Slc7a2^+/+^* littermates with glucose, arginine, or a combination of the two. Basal glucagon secretion from *Slc7a2^−/−^*islets was low, even at low glucose, and arginine did not stimulate glucagon secretion from these isolated islets (Fig. 2G). This appears to be due to a secretory defect in *Slc7a2^−/−^* α cells as glucagon content was similar in *Slc7a2^+/+^* and *Slc7a2^−/−^* islets (Supplemental Fig. 1G) and membrane depolarization with KCl failed to stimulate glucagon secretion as it robustly did in *Slc7a2^+/+^* islets (Fig 2G). Arginine-stimulated insulin secretion was impaired in the *Slc7a2^−/−^* islets (Fig. 2H), again, with no difference in insulin content (Supplemental Fig. 1H). KCl-stimulated and 2^nd^ phase glucose-stimulated insulin secretion was also impaired in *Slc7a2^−/−^* islets, but not absent. Taken together, these data indicate an islet autonomous secretory defect in *Slc7a2^−/−^* α and β cells.

### *Slc7a2^−/−^* mice have normal islet morphology and endocrine cell mass and islet-specific gene expression

*Slc7a2^−/−^* mice have elevated SLC7A2 transported amino acids, (e.g. arginine, lysine, and ornithine) (21). While these amino acids showed the greatest increases in *Slc7a2^−/−^* mice, we observed modest elevations in several other serum amino acids, including glutamine (Supp. Fig. 1F). To assess whether this slight elevation of amino acids, as compared with that of mice with interrupted glucagon signaling, is associated with increased α cell mass in the *Slc7a2^−/−^* mice, we performed immunofluorescence staining on pancreas sections from these mice. Islets from *Slc7a2^−/−^* mice have normal islet architecture and morphology (Fig. 3A-B) and no difference in α, β or δ cell mass (Fig. 3C-F) similar to no difference being observed in the glucagon or insulin content of isolated islets (Supplemental Fig. 1G-H).

**Figure 3.**
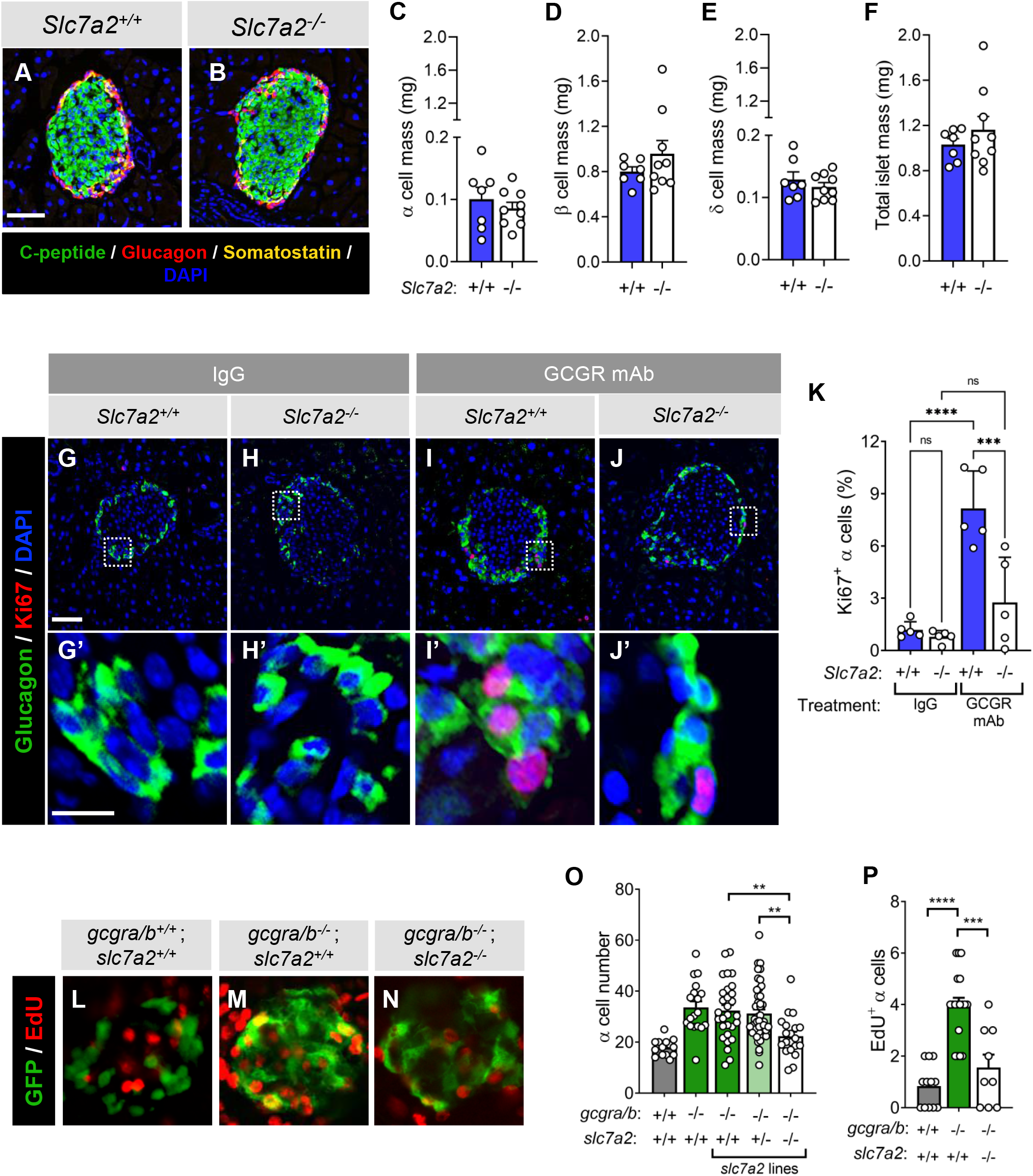
SLC7A2 is required for α cell proliferation in response to interrupted glucagon signaling. (**A-B**) Representative immunostaining for C-peptide, glucagon, and somatostatin from *Slc7a2^+/+^* and *Slc7a2^−/−^* mouse pancreas (scale bar = 50mm). (**C-E**) Mass analysis for α, β and δ cells and **(F)** total islet mass from *Slc7a2^+/+^* (n=7) and *Slc7a2^−/−^* (n=9) mice. (**G-J**) Representative images of islets from *Slc7a2^+/+^*and *Slc7a2^−/−^* mouse pancreas after two weeks treatment with GCGR mAb or control IgG (scale bar = 50µm; inset (**G’-J’**) scale bar = 10µm). (**K**) Quantification of α cell proliferation as determined by percent Ki67+/Gcg+ cells per total Gcg+ cells in *Slc7a2^+/+^* and *Slc7a2^−/−^* mouse islets after two weeks treatment with GCGR mAb or control IgG (n=5 each). (**L-N**) Representative images of day five post-fertilization islets from *Tg(gcga:EGFP)* zebrafish, α cell-specific EGFP, with CRISPR/Cas9-induced loss of glucagon receptors (*gcgra/b*) and/or *slc7a2* stained for EdU to assess proliferation (scale bar = 20µm). (**O**) Quantification of total α cell numbers (n=14, 18, 29, 46, and 20, respectively for each genotype) and (**P**) quantification of EdU positive α cells from zebrafish islets (n=12, 18, and 9, respectively for each genotype).

### Interrupted glucagon signaling-stimulated α cell proliferation requires *Slc7a2* expression

To investigate the role of SLC7A2 in α cell proliferation, we used complimentary zebrafish and mouse models of interrupted glucagon signaling where SLC7A2 expression was targeted. We previously described a *gcgr* knockout zebrafish model with increased α cell proliferation and numbers (12). Deletion of *slc7a2* by CRISPR targeting in *gcgr* null zebrafish decreased the total number of α cells and EdU-positive α cells at day five post-fertilization to wildtype fish levels, indicating that Slc7a2 is necessary for α cell proliferation in response to interrupted glucagon signaling in the developing zebrafish islet (Fig. 3L-P).

In a second model, treatment of mice with a monoclonal antibody targeting the glucagon receptor (GCGR mAb) interrupted glucagon signaling and stimulated α cell proliferation as previously shown (4, 9, 10). In *Slc7a2^+/+^*mice, two weeks treatment with GCGR mAb resulted in a greater than 6.8-fold increase in the percentage of Ki67-positive α cells (Fig 3G-H, K). *Slc7a2^−/−^* mice, treated in the same way, showed a 66% reduction in α cell proliferation (Fig 3J and K). This stimulated α cell proliferation has been mechanistically linked to mTOR signaling and increased phosphorylation of ribosomal protein S6 (phospho-S6), a target of mTOR activity (10). We observed phospho-S6 staining in α cells of Gcgr mAb treated *Slc7a2^−/−^* mice similar to IgG-treated mice indicating decreased mTOR signaling in response to interrupted glucagon signaling in *Slc7a2^−/−^*α cells (Fig 4). Together these studies in zebrafish and mouse models of interrupted glucagon signaling indicate that SLC7A2 is necessary for α cell proliferation, and that this proliferative response is mechanistically linked to downstream mTOR signaling. Interrupted glucagon signaling does not influence the pattern of secretory profile in response to stimuli; rather the genotype predicts the secretory response including no response to the glucose/arginine bolus in *Slc7a2^−/−^* mice (Supplemental Fig. 3). However, the overall amount of glucagon was increased 5-fold in all genotypes by GCGR mAb treatment versus IgG-treated littermates suggesting that glucagon secretion in *Slc7a2^−/−^* mice is blunted but not completely lost in response to other amino acids that are not transported by SLC7A2.

**Figure 4.**
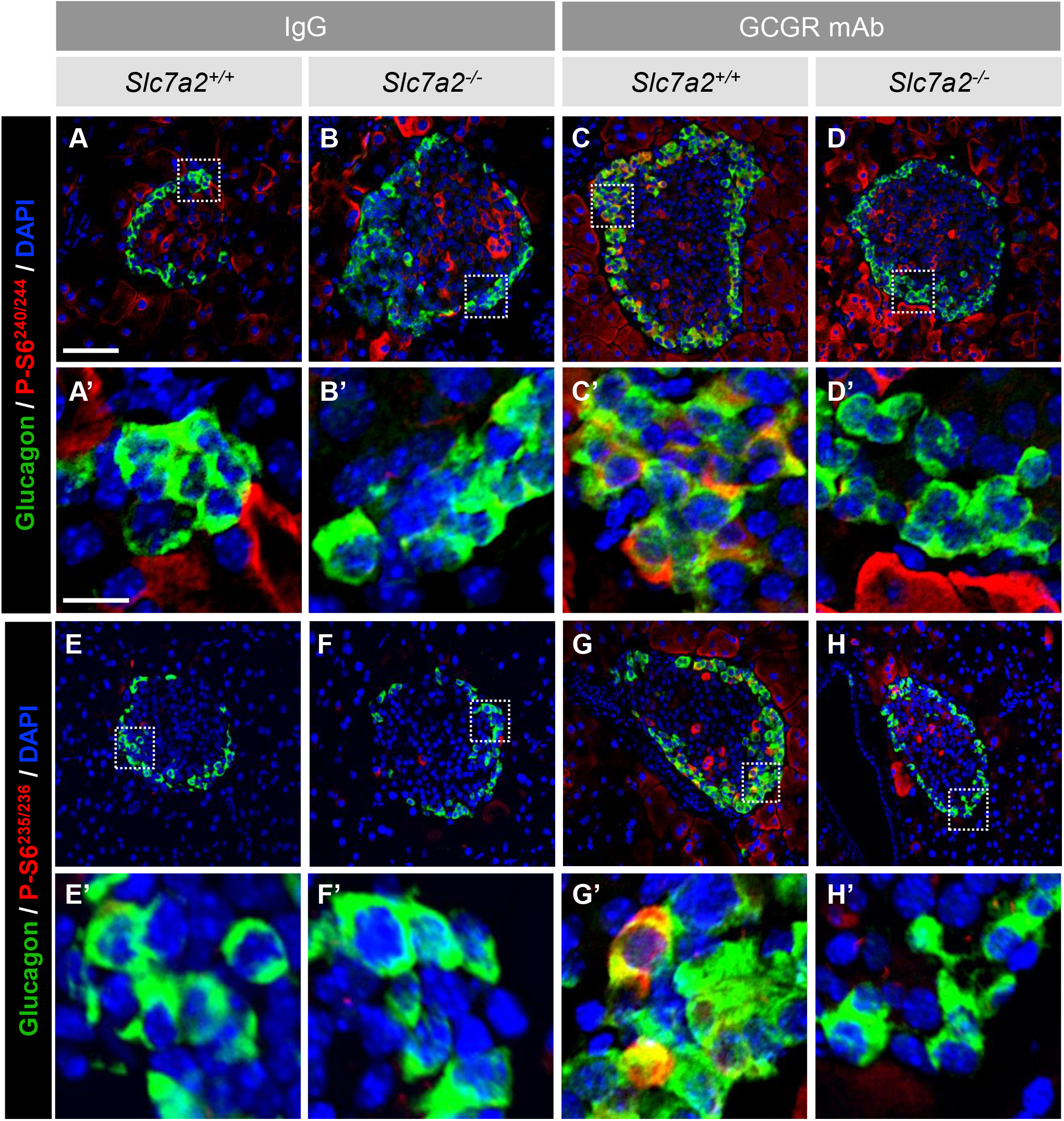
Arginine stimulation of α cell proliferation is mTOR-dependent. Immunostaining of GCGR mAb and IgG treated *Slc7a2^+/+^* and *Slc7a2^−/−^* mouse islets for glucagon and phospho-ribosomal protein S6 (P-S6^240/244^) (**A-D**) and (P-S6^235/236^) (**E-H**), indicating active mTOR signaling (scale bar = 50µm; inset (**A’-H’**) scale bar = 10µm).

### SLC7A2-dependence of stimulated α cell proliferation is islet cell autonomous

To evaluate whether reduced α cell proliferation in *Slc7a2^−/−^*mice is an islet autonomous or an extra-islet effect, we first evaluated SLC7A2-dependent proliferation in clonal αTC1-6 shRNA cell lines. Two separately selected monoclonal *Slc7a2* shRNA lines showed decreased SLC7A2 protein as compared to a non-targeting (scrambled) shRNA expressing line (Fig. 5A, right). Evaluation of growth of these monoclonal lines over eight days demonstrated that SLC7A2 loss reduced proliferation in these cells with cell numbers approximately 65% lower in *Slc7a2* shRNA lines (Fig. 5A). These data, when combined with the findings presented in Supplemental Figure 1B indicate that arginine and SLC7A2 are necessary for growth of the αTC1-6 mouse α cell line.

**Figure 5.**
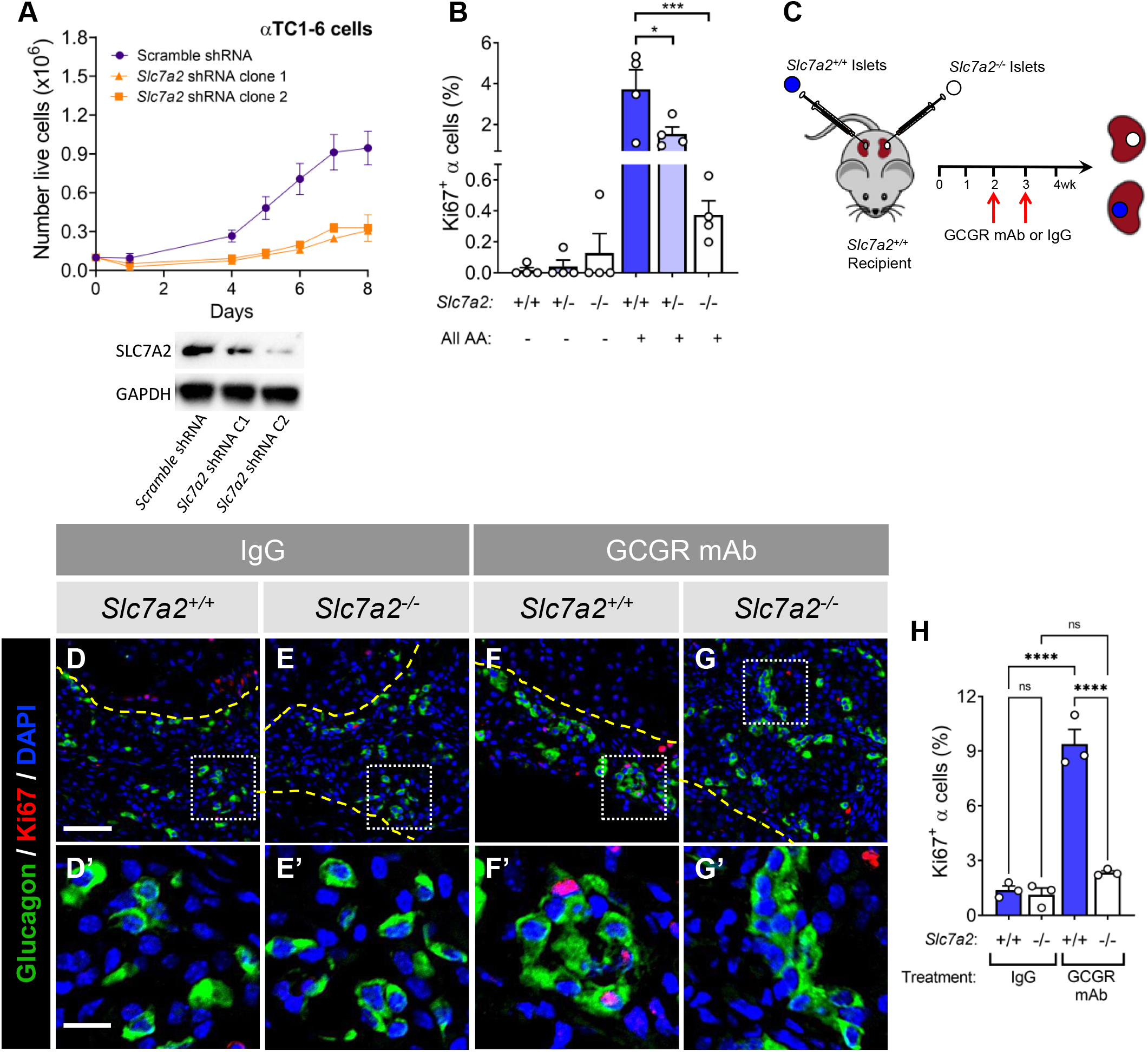
SLC7A2-dependent stimulated α cell proliferation is islet autonomous. (**A**) Cell growth over time of αTC1-6 cultured cells expressing *Slc7a2* shRNA (2 clones) or non-targeting (scrambled) control shRNA (n=4 each). Immunoblot (right) shows decreased SLC7A2 protein in the *Slc7a2* shRNA lines. (**B**) *Ex vivo* α cell proliferation from isolated mouse islets cultured in low (-) and high (+) amino acid containing medium as determined by percent Ki67+/Gcg+ cells per total Gcg+ cells in islets isolated from *Slc7a2^+/+^*, *Slc7a2^+/−^* and *Slc7a2^−/−^* mice (n=4 each). (**C**) Schematic of kidney capsule transplantation of *Slc7a2^−/−^* and *Slc7a2^+/+^* islets into *Slc7a2^+/+^* recipient mouse followed by glucagon receptor monoclonal antibody treatment. (**D-G**) Representative images of *Slc7a2^+/+^* and *Slc7a2^−/−^* islet grafts from *Slc7a2^+/+^* kidney capsules after two weeks treatment with glucagon receptor monoclonal antibody (GCGR mAb) or control IgG (scale bar = 50µm; inset (**D’-G’**) scale bar = 10µm). Dashed yellow lines indicate kidney-graft boundary. (**E**) Percent α cell proliferation from *Slc7a2^+/+^* and *Slc7a2^−/−^* islets transplanted into *Slc7a2^+/+^* kidney capsule and treated with GCGR mAb or control IgG (n=8 transplant recipients, n=4 per treatment group).

Isolated islets from *Slc7a2^−/−^* mice had a 90% reduction in α cell proliferation when cultured in high amino acids (Fig. 5B). Interestingly, proliferation of *Slc7a2^+/−^* α cells was approximately 60% lower than *Slc7a2^+/+^*, suggesting that the level of *Slc7a2* expression may be rate limiting for proliferative response, but not the glucagon secretory response, to arginine.

To test islet autonomy *in vivo*, isolated islets from *Slc7a2^−/−^*mice were transplanted into *Slc7a2^+/+^* recipients, placing the knockout islets into the physiological environment of a wild type mouse. Each recipient mouse also had *Slc7a2^+/+^*islets transplanted into the contralateral kidney to serve as a control for stimulated α cell proliferation. After engraftment, recipient mice were treated weekly with either the GCGR mAb or the control IgG for an additional two weeks to investigate the consequence of interrupted glucagon signaling (Fig. 5C). Immunostaining analysis of kidney grafts indicated that interrupted glucagon signaling increased α cell proliferation 4.5-fold in the *Slc7a2^+/+^* islet grafts but α cells in *Slc7a2^−/−^* grafts did not proliferate above control, IgG treatment (Fig. 5D-G and H). Thus, these *in vitro* and *in vivo* data together indicate that SLC7A2-dependence of stimulated α cell proliferation is likely a result of SLC7A2 function in α cells.

### Stimulated *Slc38a5* expression during interrupted glucagon signaling is SLC7A2-dependent

Previous studies showed that interrupted glucagon signaling stimulates expression of *Slc38a5*, a neutral amino acid/glutamine transporter, in mouse and zebrafish α cells, and that this is partially required for α cell proliferation in an elevated amino acid environment associated with interrupted glucagon signaling in liver (9, 10). To test whether *Slc38a5* expression was SLC7A2-dependent, we evaluated the presence of SLC38A5 in α cells of GCGR mAb treated *Slc7a2^−/−^* mice and *Slc7a2^+/+^* littermates by immunofluorescence (Fig. 6A-D). As expected, SLC38A5 was detected in greater than 60% of GCGR mAb-treated *Slc7a2^+/+^*α cells versus 10% in IgG-treated littermates. Interestingly, less than 20% of *Slc7a2^−/−^* α cells showed detectable SLC38A5 similar to IgG treated mice of either genotype (Fig. 6E). Isolated *Slc7a2^−/−^* islets transplanted into the kidney capsules of *Slc7a2^+/+^* mice subsequently treated with GCGR mAb also lacked this stimulated SLC38A5 expression (Fig 6F-I). Isolated islets from *Slc7a2^−/−^* mice treated with GCGR mAb also have impaired induction of *Slc38a5* gene expression (Fig 6J).

**Figure 6.**
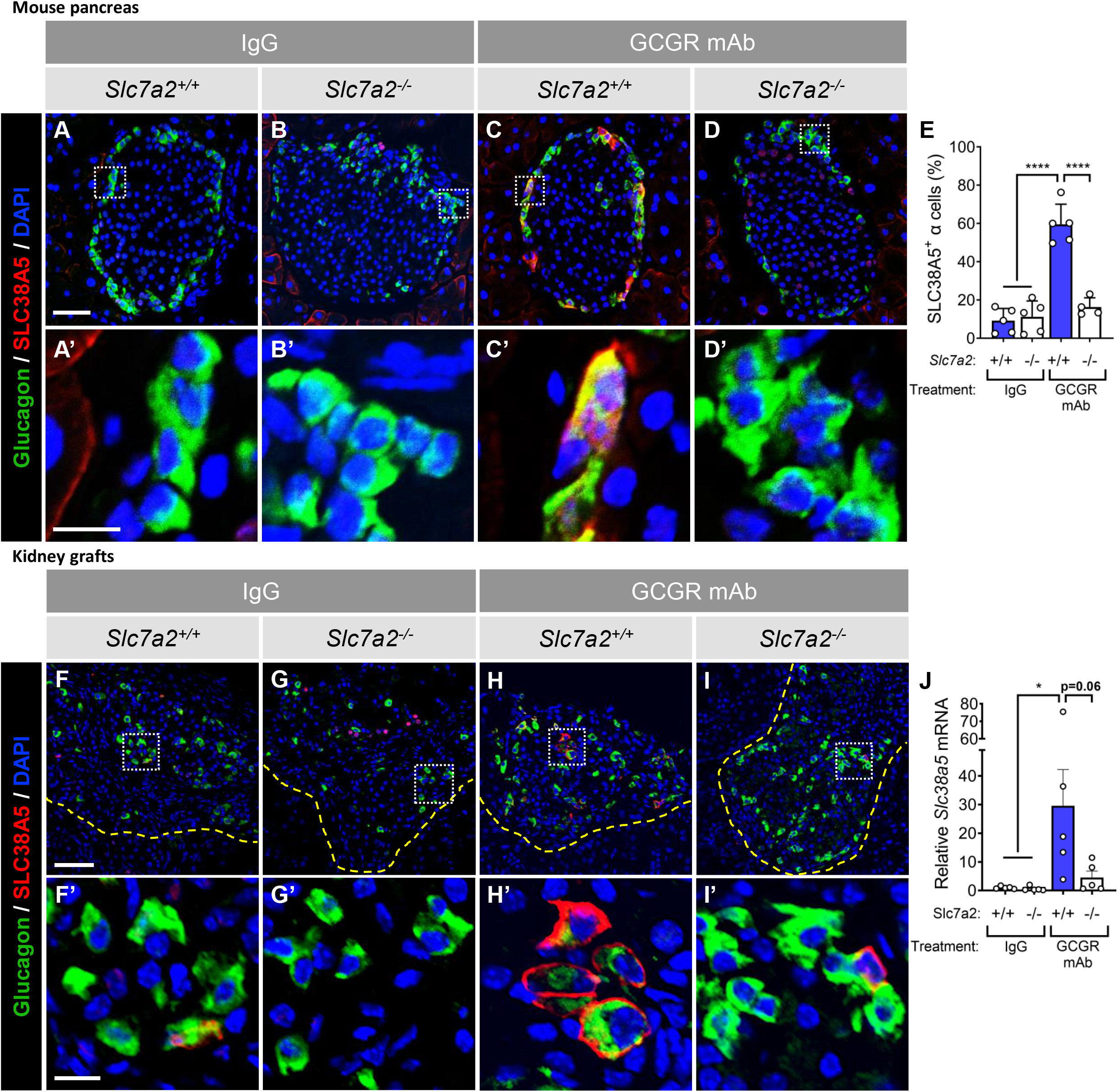
SLC7A2 is required for upregulation of *Slc38a5* expression during interrupted glucagon signaling. (**A-D**) Representative images of islets from *Slc7a2^+/+^* and *Slc7a2^−/−^* mouse pancreas stained for glucagon (green) and SLC38A5 (red) after two weeks treatment with GCGR mAb or control IgG (scale bar = 50µm; inset (**A’-D’**) scale bar = 10µm). (**E**) Quantification of SLC38A5 expression in α cells by percent SLC38A5+/GCG+ cells per total Gcg+ cells (% SLC38A5^+^ α cells) in *Slc7a2^+/+^*and *Slc7a2^−/−^* mouse islets after injection with GCGR mAb or control IgG (n=4-5). (**F-I**) Representative images of *Slc7a2^+/+^*and *Slc7a2^−/−^* islet grafts from *Slc7a2^+/+^* kidney capsules after two weeks treatment with GCGR mAb or control IgG and stained for glucagon and SLC38A5 (scale bar = 50µm; inset (**F’-I’**) scale bar = 10µm). Dashed yellow lines indicate kidney-graft boundary.

### *SLC7A2* is associated with HbA1C levels in humans

To ask if *SLC7A2* contributes to nutrient homeostasis in humans, we investigated if the *SLC7A2* gene locus is associated with diabetes-related phenotypes in human GWAS. Two single nucleotide polymorphisms (SNPs) have been identified within the first intron of *SLC7A2* that are associated with HbA1c levels (Fig 7A). rs142010226 (chr8, 17367112:A/G) and rs2517232 (chr 8, 17367421:A/G) are both strongly associated (P < 10^-15^) with HbA1c in the EXTEND human cohort, which consists of 1,395 diabetic and 5,764 non-diabetic individuals of European ancestry which we identified via datamining the T2D Knowledge Portal (27). Further datamining of published ATAC-seq and ChIP-seq datasets from human islets (Fig 7B) show that both SNPs occur within ∼1 kb of binding sites for MAFB and FOXA2, two transcription factors (28, 29) that have previously shown to be critical for α cell gene expression (30–33). In summary, these findings are consistent with a role for *SLC7A2* in glucose homeostasis in humans and suggest that genetic variants associated with HbA1c may influence MAFB and/or FOXA2-depndent regulation of *SLC7A2* expression in islets.

**Figure 7.**
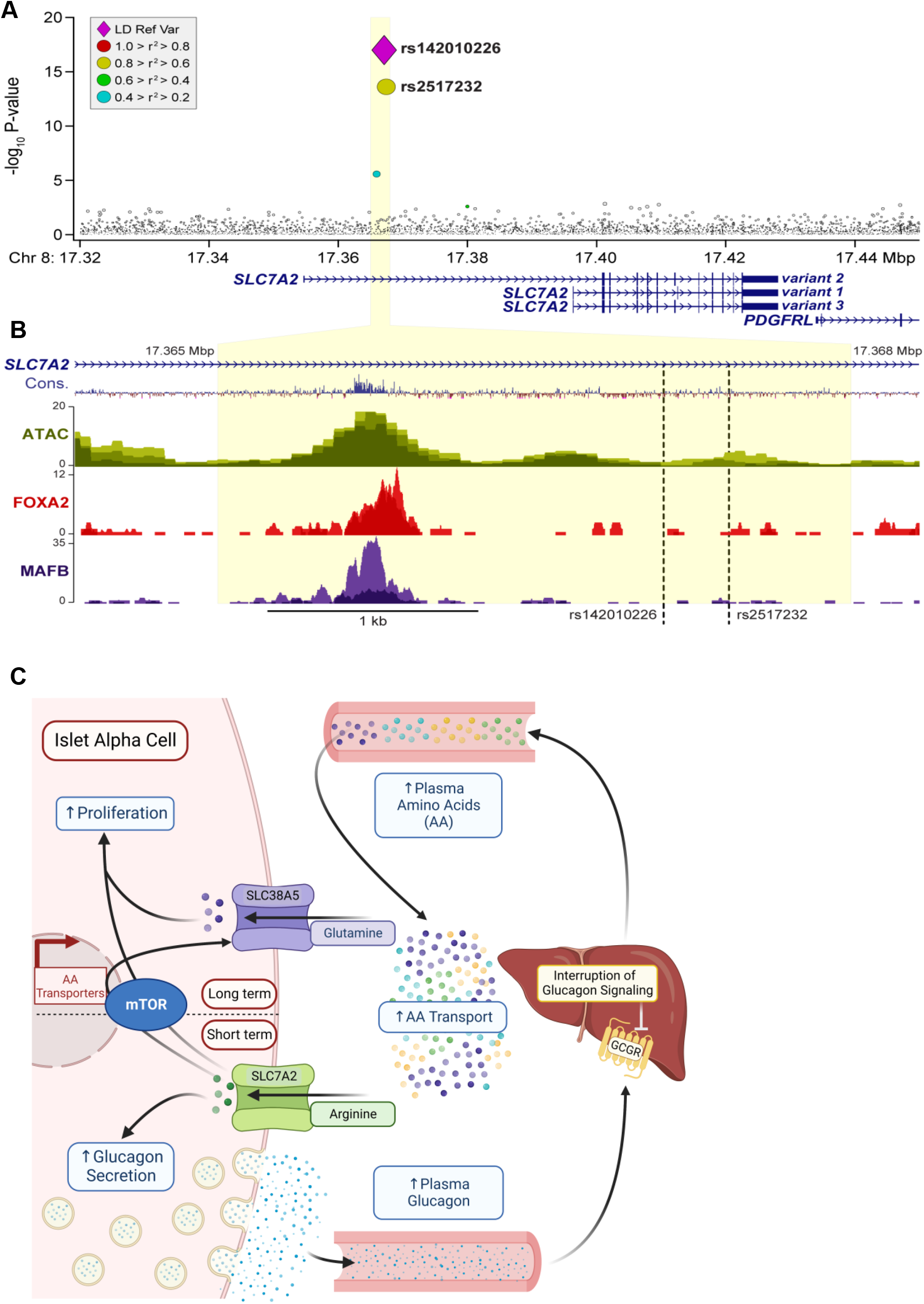
Model of α cell response to elevated amino acids during interrupted glucagon signaling. (**A**) Single nucleotide polymorphisms (SNPs) significantly associated with the human *SLC7A2* gene locus with Hemoglobin A1C (HbA1C) levels are represented by a violet diamond (rs142010226) and a yellow (rs2517232) circle. Gray circles represent SNPs not significantly associated with HbA1C or for which there is no associated data, respectively. Yellow shading indicates the region identified in **(B)** ATAC-Seq analyses (from Pasquali et al (28)) of the *SLC7A2* locus in human islets with highly conserved regions (blue-Cons), ATAC peaks in green, FOXA2 binding in red, and MAFB binding in purple. **(C)** SLC7A2 is required for arginine-stimulated glucagon secretion and α cell proliferation. Arginine transported into the α cell through SLC7A2 initially stimulates glucagon secretion (Short term). The build-up of intracellular arginine ultimately stimulates mTOR-dependent *Slc38a5* expression (Long term). Transport of glutamine through SLC38A5 stimulates α cell proliferation as described previously (9, 10).

## Discussion

Overall, these data suggest that arginine transport through SLC7A2 is required for the induced gene expression of the glutamine transport SLC38A5 observed during interrupted glucagon signaling (Fig. 7C). Arginine is a major substrate for SLC7A2 transport and a known potent secretagogue for both pancreatic islet α and β cells. We found that SLC7A2, one of the most highly expressed amino acid transporter genes in α cells in mice and humans, plays a central role in islet function. Loss of *Slc7a2* expression in mice resulted in impaired arginine-stimulated glucagon and insulin secretion *in vivo* and *ex vivo. Slc7a2* expression is also required for amino-acid stimulated α cell proliferation in zebrafish and mice. Together, our data indicate that SLC7A2 is the primary arginine transporter in islet cells regulating arginine’s effects on hormone secretion. Our work also provides strong evidence that arginine stimulation of hormone secretion works by an alternative mechanism than what has been previously suggested (35). These studies directly link SLC7A2 function with the previous observation that glutamine transport through SLC38A5 is necessary for α cell proliferation by demonstrating that arginine transport through SLC7A2 is required for mTOR activation and upregulation of *Slc38a5* expression (see model Fig 7C). Finally, we show that SNPs in the SLC7A2 gene are associated with HbA1c, a common clinical value used to evaluate glycemic status. These newly identified roles for SLC7A2 in α cells described here support its designation as an α cell “signature gene” (34).

Previous studies suggested that arginine transport results in membrane depolarization due to the cationic properties of arginine (37). Yet, glucagon secretion from *Slc7a2^−/−^* islets was not stimulated in perifusion experiments even with the strong depolarizing agent KCl while glucagon content and overall α cell mass were not different (Fig 3). This suggests that impaired hormone secretion when SLC7A2 expression is lost represents a fundamental defect in secretory mechanisms or machinery versus secretory capacity and not simply perturbations in membrane polarity. This was surprising since we predicted that KCl stimulation would remain intact in *Slc7a2^−/−^* islets based on previous studies. Whole islet transcriptomics studies in *Slc7a2^−/−^* and *Slc7a2^+/+^* islets did not reveal any insight into a possible mechanism (e.g. changes in ion channel or vesicle machinery expression; Supp. Fig 2). Therefore, further study is needed to determine how arginine and SLC7A2 promote secretion.

Our work connects arginine signaling to glutamine signaling by showing *Slc7a2^−/−^* mice lack the increased expression of the glutamine transporter, *Slc38a5*, seen in *Slc7a2^+/+^* α cells, and described in previous studies (9, 10) when glucagon signaling is interrupted. Cancer cells also upregulate amino acid transport and metabolic pathways to facilitate their growth but under nutrient limiting conditions. In cancer cells, arginine-dependent regulation of cytosolic CASTOR1 and lysosomal SLC38A9 and TM4SF5 proteins promote mTOR activation leading to proliferation (47). Whether or not these pathways for arginine-stimulated proliferation are conserved in non-transformed cells including α cells is unclear. In contrast to the present study, SLC7A2 expression is negatively correlated with proliferation in breast, colon, hepatocellular carcinoma, and ovarian cancers (48–51). SLC7A2 is highly expressed in multiple normal tissues including liver, breast, skeletal muscle, and islets. Thus, the role of SLC7A2 in cell proliferation and physiology in general merits cell type-specific targeting for further study.

*SLC7A2/Slc7a2* is alternatively spliced at exon 7 yielding two protein isoforms CAT2A and CAT2B (17). While both exons appear to be expressed in α and β cells, α cells predominately express the exon the 2^nd^ exon 7 associated with SLC7A2A (CAT2A) isoform and β cells prefer the 1^st^ exon 7 associated with SLC7A2B (CAT2B). This suggests that α cells have higher levels of CAT2A a low affinity (Km 2-5mM), high-capacity arginine transporter. This difference, in addition to overall higher expression, might grant α cells the ability to sense rising levels of arginine better than other islet cells. Interestingly, *SLC7A2* is downregulated in both T1D and T2D human α cells and may reflect observed downregulation of *MAFB*/MAFB under both conditions (22, 38-40) which we propose regulates *SLC7A2* expression via binding to the first intron of *SLC7A2*. The idea that MAFB regulates *Slc7a2*/*SLC7A2* gene expression is supported by gene knockout studies in mice where loss of *Mafb* during either pancreatic islet development or postnatal period (5 weeks of age) result in significantly reduced *Slc7a2* expression and arginine-stimulated glucagon secretion (33).

Our analysis of the association to HbA1c levels in the EXTEND GWAS data set suggests that individuals with either of two SNPs within the first intron of SLC7A2 have reduced HbA1c levels (i.e., negative β values). It is unclear if this association is due to glycemia as only a third of variance in HbA1c levels in non-diabetic individuals is due to factors such as glycemia and body mass index (41). That these SNPs are proximal to binding sites for MAFB and FOXA2 in human islets, suggests they may influence their binding and subsequent regulation of *SLC7A2* expression. In mouse islets *Slc7a2* expression is positively correlated with the expression of *Foxa2* and *Mafb* (42). Attie and colleagues also show that genetics exerts a strong influence on SLC7A2 protein levels in pancreatic islets from different mouse strains (43). Their proteomic survey showed that islets from CAST mice have ∼6 and ∼11-fold lower SLC7A2 protein levels than islets from B6 or 129 mice, respectively. In our mouse studies, loss of SLC7A2 expression results in higher blood glucose in response to an arginine and glucose bolus. It is possible that these SNPs in human enhance FOXA2 and/or MAFB binding, leading to increased expression of SLC7A2 and thus lower circulating blood glucose. However, it is unclear where SLC7A2 expression is driving such associations since MAFB is expressed in both human α and β cells and FOXA2 is expressed in both α cells and liver. Therefore, future studies with cell type specific SLC7A2 targeting are warranted. These studies suggest that differences in SLC7A2 levels may underlie differential responsivity to arginine-induced glucagon and insulin secretion.

Despite associations of SNPs in *SLC7A2* with improved long-term glucose homeostasis in humans, *Slc7a2^−/−^* mice had normal fasting glucose and glucose tolerance when assessed by acute intraperitoneal (IP) glucose injection. Rather our study supports that arginine regulation of hormone secretion itself may be the driver behind the association and not glucose disposal per se. Furthermore, there remains uncertainty as to whether the observed insulin secretion defect is β cell autonomous or the result of impaired paracrine signaling from α cells. Interestingly, early studies with the perfused rat pancreas showed that arginine rapidly and potently stimulates glucagon secretion under no glucose conditions whereas insulin secretion is slow and tonic requiring several minutes for appreciable insulin secretion suggesting that SLC7A2 transporter levels limit the response by islet cells to arginine (44) similar to the intermediate effect of an arginine bolus on insulin levels in *Slc7a2^+/−^* mice versus *Slc7a2^−/−^*mice observed in this study. However, under high glucose conditions both glucagon and insulin are rapidly and potently secreted in response to arginine. This argues that other factors than SLC7A2 transporter levels may limit β cell responses to arginine. The glucose-dependency observed in this study is similar to responses observed with incretin-stimulated insulin secretion where incretins have little effect on insulin secretion under low or euglycemic conditions. More recently, Capozzi, Campbell, and colleagues demonstrated that intra-islet glucagon signaling through the GLP-1 receptor is required for amino acid-stimulated insulin secretion, including arginine (45). Similarly, Svendsen, Holst, and colleagues concluded that both glucagon and GLP-1 receptor signaling in β cells were required for arginine-stimulated insulin secretion (46). Interestingly, isolated islets from mice where α cells had been selectively ablated had dramatically reduced arginine-stimulated insulin secretion (46). We observed a modest decrease in glucose-stimulated insulin secretion in *Slc7a2^−/−^* islets while arginine-stimulated insulin secretion was profoundly perturbed in addition to a loss of glucagon secretion. Therefore, we predict that much of arginine’s effect on β cells might be paracrine in nature mediated via proglucagon-derived peptides linked to the α cell’s robust expression of SLC7A2. To understand whether the loss of arginine-stimulated insulin secretion in *Slc7a2^−/−^*mice results from the intrinsic loss of SLC7A2 from β cells or whether it is the result of the observed loss of glucagon secretion, α and β cell-specific *Slc7a2* knockout mice would be necessary.

It was surprising that no difference in α cell mass were observed in untreated *Slc7a2^−/−^* mice since SLC7A2 was required for adaptive growth in response to hyperaminoacidemia and glucagon secretion was nearly absent under our conditions. This suggests that other mechanisms might also determine baseline α cell mass. However, we predict that there is sufficient glucagon signal in the liver in these mice to not fully activate the liver-α cell axis as with pharmacological or genetic targeting glucagon receptors. This hypothesis is supported by the observation that while *Slc7a2^−/−^* mice have mild hyperaminoacidemia, blood levels of amino acids such as glutamine and arginine are 3-5 fold lower than levels observed in mice with interrupted glucagon signaling (9–11). Expression of *Slc38a5* is greatly increased in α cells during interrupted glucagon signaling and it is the only amino acid transporter that responds as such. Interestingly, *Slc38a5^−/−^* mice only showed 50% lower α cell proliferation during interrupted glucagon signaling than wild type mice indicating that other amino acids or transporters are required for the α cell proliferative response (9). These observations led us to hypothesize that other amino acids may play an essential role in α cell proliferation.

Arginine plays several critical physiological roles including controlling vasodilation through the synthesis of nitric oxide, immune function, and removing ammonia from the body via production of urea. Glucagon facilitates ureagenesis by both transcriptional and post-translational control of the urea cycle in liver. Conversely, several amino acids including arginine potently stimulate glucagon secretion. Together, these events form an endocrine feedback loop called the liver-α cell axis where α cells play a critical role in the sensing of circulating amino acids (Fig 7) (1, 36). When arginine levels rise acutely (e.g., minutes to hours) such as with a protein meal, arginine transport via SLC7A2 leads to glucagon secretion. Similarly, chronic hyperargininemia (e.g., days or longer), such as with interruption of glucagon signaling in liver, and prolonged arginine accumulation via SLC7A2 leads to mTOR activation in α cells and activation of gene expression (including other amino acid/glutamine transporters such as *Slc38a5*) that facilitate cell proliferation.

The discovery that interrupted glucagon signaling stimulates human α cell proliferation (4, 14) and that human α cells can transdifferentiate into β cells (52) holds promise for the use of glucagon receptor antagonists for the reestablishment of β cell mass after diabetic loss as has been recently demonstrated in mice (4). Safely expanding α cells could also represent the first step in restoring β cell mass through α-to-β cell transdifferentiation. Further studies are needed to address the molecular mechanisms by which arginine stimulates glucagon secretion, arginine interacts with the mTOR pathway in α cells to stimulate *Slc38a5* expression and proliferation, whether transport of other amino acids is required for these responses, and whether these mechanisms are conserved in human islets.

### Research Design and Methods

#### Mouse studies

All mouse studies were performed at Vanderbilt University Medical Center and approved by the Institutional Animal Care and Use Committee. Mice were housed on a 12:12 hour light:dark cycle with *ad libtum* access to standard rodent chow (unless indicated for fasting purposes) and water. *Slc7a2^−/−^*mice (Jackson labs, B6.129S7-*Slc7a2^tm1Clm^*/LellJ) and *Slc7a2^+/−^*and *Slc7a2^+/+^* littermates from heterozygous crosses were used for all mouse experiments. Both male and female mice were combined in these studies as there were no differences were observed between sexes. To interrupt glucagon signaling, mice were treated weekly with 10mg/kg of a humanized monoclonal antibody targeting the glucagon receptor (GCGR mAb “Ab-4”) intraperitoneally once a week for 2 weeks (53).

For contralateral islet transplantations, 100-150 islets isolated from 14-16-week-old *Slc7a2^−/−^* donor mice were transplanted under the left kidney capsule and an equivalent number of *Slc7a2^+/+^* donor islets were transplanted under the right kidney capsule of a *Slc7a2^+/+^* recipient mouse. Two weeks after transplantation, transplant recipients were given two weekly treatments of GCGR mAb as described in Fig 5C and above. After two weeks of GCGR mAb treatment, kidneys containing grafts were retrieved, bisected at the grafts, fixed in 4% paraformaldehyde, embedded in OCT, and stored at −80°C until sectioning (14, 54).

#### Zebrafish studies

The role of Slc7a2 in proliferation of α cells in Zebrafish (*Danio rerio*) was studied using a previously described (12) glucagon receptor double-knockout line expressing GFP under the control of the glucagon promoter to identify α cells (*Tg(gcga:GFP)*;*gcgra/b^−/−^*). Using CRISPR/Cas9 technology, a 62bp deletion was created in *slc7a2* (using the sgRNA GgGTAAGCGCCAGTCGCCAG and the PAM TGG for targeting). We assessed total α cells by counting GFP^+^ cells in the islets of five days post-fertilization (dpf) fish as described previously (12). To identify proliferating α cells, embryos were incubated with 1 mmol/l 5-ethynyl-2-deoxyuridine (EdU) at 4 dpf and chased for 24 h. EdU was detected according to published protocols (12) and using the Click-iT EdU Alexa Fluor 594 Imaging Kit (C10339; Invitrogen). All images were collected using a Zeiss LSM880 confocal microscope (Carl Zeiss, Jena, Germany).

#### Tissue collections from Mice

After treatment periods, pancreata were collected. Histology samples were fixed in 4% paraformaldehyde, embedded in OCT (Fisher Scientific) and stored at −80°C until use. For transplantation, islet RNA and *ex vivo* proliferation experiments, islets were isolated by intraductal infusion of collagenase P and histopaque gradient separation.

#### Stimulated glucagon and insulin secretion *in vivo*

To assess the role of SLC7A2 in glucagon and insulin secretion, *Slc7a2^+/+^*, *Slc7a2^+/−^* and *Slc7a2^−/−^* mice were fasted for 6 hours and then injected intraperitoneally with glucose, arginine, or both to final concentrations of 2g / kg body weight each. Blood was collected retroorbitally before injection (Fasting) and 15 minutes after injection (Stimulated). Blood glucose was measured with a hand-held glucometer (Accu-Check Aviva) and remaining whole blood was spun, serum collected into separate tubes and stored at −80°C for glucagon and insulin analyses.

Serum hormones were analyzed in the Vanderbilt Hormone and Analytical Services core. Serum glucagon was analyzed in a two-site enzyme sandwich ELISA (Mercodia). Serum insulin was analyzed by dual antibody radioimmunoassay and counted in a Packard Gamma counter.

#### Immunofluorescence staining and image analysis

To compare islet cell masses in untreated Slc7a2^+/+^ and Slc7a2^−/−^ mice, pancreas from 14-16-week-old mice were prepared for histology as described above. Whole mouse pancreas was sectioned on a cryostat at a thickness of 8μm per section. The full depth of the pancreas was sectioned by seven repeated steps of sectioning away 150μm and collecting 10 sections onto slides. In this way 7 depths of 150μm were collected from each pancreas. For islet cell mass analysis, one section from each of the seven depths was stained for C-peptide (Invitrogen, PA-85595) to mark β cells, glucagon (LSBio, LS-C202759) to mark α cells and somatostatin (Santa Cruz, sc-7819) to mark δ cells. Whole pancreatic sections were imaged on Scanscope FL System (Aperio Technologies) and islet cell areas were analyzed using Halo image analysis software (Indica Labs). Total pancreatic islet cell masses were calculated as described previously (Golson 2014). Briefly, islet cell areas from each of seven depths were normalized to total pancreas section area, areas from each of the seven sections were summed and the sum multiplied by total pancreas mass to achieve an estimate of the percent of total mass.

For α cell proliferation analysis, pancreas form mice treated for two weeks with GCGR mAb were sectioned and immunostained for the proliferation marker, Ki67 (Abcam, ab15580), and glucagon to mark α cells. Whole sections were imaged on the Scanscope and analyzed on Halo software. Percent α cell proliferation was calculated from at least 1000 α cells per animal by dividing Ki76^+^/glucagon^+^ cells by total glucagon^+^ cells. Similarly, percent Slc38a5^+^ α cells were calculated by staining for Slc38a5 (Santa Cruz, sc-50682) and glucagon, imaging and analyzing as above. Amino acid-stimulated α cell proliferation has been shown to be mTOR-dependent. We verified this mTOR-dependence by staining for a target of mTOR1 activity, phosphorylation of ribosomal protein S6 (Cell Signaling: pS6^235/236^, 4858S; pS6^240/244^, 5364S) and glucagon.

Kidney grafts were sectioned at 5μm per section and approximately 20 sections were collected for each graft. These sections were immunostained for glucagon and Ki67 to assess percent α cell proliferation from at least 500 α cells per graft. Grafts were also stained for glucagon and SLC38A5 to assess α cell-specific expression of the transporter during interrupted glucagon signaling.

#### *Ex vivo* α cell proliferation

*Ex vivo* α cell proliferation was assessed as described previously (10). Briefly, islets isolated from *Slc7a2^−/−^*, *Slc7a2^+/−^*and *Slc7a2^+/+^* mice were cultured in DMEM-based medium with high or low amino acid concentrations, based on amino acid levels in *Gcgr^−/−^* and *Gcgr^+/+^*mouse serum, respectively (See Supplemental Table 1 for amino acid concentrations in islet culture media), for 72 hours. After culture, islets were washed in 2mM EDTA and dispersed with 0.025% Trypsin at 37°C for 10 minutes with mixing. Dispersed islet cells were recovered by centrifugation in RPMI media containing 5.6mM glucose, 10% FBS, and 1% Penicillin/Streptomycin. The resulting cell pellet was resuspended in medium and centrifuged onto a glass slide using a Cytospin 4 (Thermo Scientific, Waltham, MA) centrifuge. Slides were air-dried for 30 minutes, then stored at −80C until use. For staining, slides were thawed, immediately fixed in 4% paraformaldehyde and immunostained for glucagon, to mark α cells, and Ki67, to mark proliferating cells. Slides were imaged using a Leica Microsystems Epifluorescent Microscope DM1 6000B. The percent of proliferating α cells was calculated based on the number of Ki67^+^/glucagon^+^ cells divided by the total number of glucagon^+^ cells using Imaris image analysis software (Oxford Instruments).

#### αTC1-6 cell culture

α TC1 clone 6 (αTC1-6) cells were obtained from ATCC (CRL-2934) and maintained in DMEM, low glucose (Gibco 11885-084) with 10% FBS, 15mM HEPES, 0.1mM non-essential amino acids, 0.02% bovine serum albumin and 1% Penicillin/Streptomycin (Supplemental Table 2). To test the roles for arginine and glutamine in growth of these cells, SILAC DMEM Flex Media (Gibco, A24939-01), which lacks arginine, glutamine and lysine, was used as base medium, prepared as above, and supplemented with lysine to produce ‘No Gln / No Arg’ control medium. The control medium was supplemented with either 0.4mM arginine (No Gln / 0.4mM Arg) or 4mM glutamine (4mM Gln / No Arg) or both (4mM Gln / 0.4mM Arg) to test the effects of glutamine and arginine on αTC1-6 cell growth. To assess cell growth, 6 well plates were seeded with 1 x 10^5^ cells/well (Day 0) in standard culture medium and allowed to normalize overnight. Cells form one well in each 6-well plate were collected and counted (Day 1) in a Countess 3 automated cell counter (Thermofisher Scientific). Media were changed in the remaining wells to experimental media described above and cells were allowed to grow for three days. Beginning on Day 4, cells from one well in each media condition were collected and counted and growth curves were established (Fig 1C).

To establish the requirement for SLC7A2 in αTC1-6 cell growth, monoclonal lines expressing *Slc7a2* shRNA or a Scrambled (non-targeting) shRNA control were established. Cells were transduced with lentivirus expressing *Slc7a2* shRNA (VectorBuilder vector no. VB170727-1121dxf) or Scrambled shRNA (VectorBuilder vector no. VB170313-1108xyf), each expressing a Puromycin resistance cassette and a GFP fluorescence marker. Transduced cells were selected with 2.5μg/mL Puromycin Dihydrochloride (Corning, 61-385-RA) to produce polyclonal lines expressing these shRNAs. Polyclonal lines were seeded at a density of one cell per well in 96-well plates and allowed to grow into single colonies in each well. Colonies were expanded and selected based on Slc7a2 expression (Fig 5A, western blot). Cell growth of these monoclonal shRNA lines was assessed as described above in basal DMEM-based medium.

#### Analysis of published RNAseq datasets

Normalized RNAseq data were obtained from the Gene Expression Omnibus (GEO) repository. Published datasets for sorted human (22, 23), mouse (24) and zebrafish (25) α and β cells were selected for comparison of transporter expression between the two cell types. Normalized expression data for all solute carriers (SLC genes) in α and β cells were sorted from highest to lowest expression and the top 50 in each cell type are given in Supplemental Table 3. The five most highly expressed cationic amino acid transporters in human α cells, sorted from highest to lowest, are shown for all three species in Fig 1.

#### Whole Islet RT-PCR

For RT-PCR analyses, *Slc7a2^+/+^* and *Slc7a2^−/−^* mice were treated with IgG or GCGR mAb or untreated. Islets were isolated as described above. RNA was isolated from whole Islets and trace DNA was removed with the RNAqueous^TM^ micro total RNA isolation kit (ThermoFisher). RNA integrity was evaluated by Agilent 2100 Bioanalyzer. cDNA was synthesized from High integrity (RIN>7) total RNA using HighCapacity cDNA Reverse Transcription Kit (Applied Biosystems) according to the manufacturer’s instructions. mRNA levels were assessed by quantitative PCR using the TaqMan assay system. Deletion of *Slc7a2* was validated by exon 2-specific quantitative real-time RT-PCR on RNA isolated from *Slc7a2^−/−^* and wild type littermate (*Slc7a2^+/+^*) islets (Supp. Fig 2B). Primers were purchased from ThermoFisher: *Actb* (internal control), Mm02619580_g1; *Slc7a2*, Mm00432032_m1; and *Slc38a5*, Mm00549967_m1.

#### *In vitro* islet perifusion

Function of isolated *Slc7a2^+/+^* and *Slc7a2^−/−^* islets was studied in a dynamic cell perifusion system at a perifusate flow rate of 1 mL/min as described previously (22, 55). Stimulus concentrations and exposure times are shown in Fig 2 traces. The effluent was collected at 3-minute intervals using an automatic fraction collector. Glucagon and insulin concentrations in each perifusion fraction and islet extracts were measured by radioimmunoassay (MilliporeSigma).

#### Statistical analysis

All data are shown with error bars indicating standard error of the mean. Data within individual experiments were compared with ordinary one-way ANOVA using Tukey correction for multiple comparisons, unless otherwise designated in the figure legend.

## Acknowledgements

**Authors Contribution** Conceptualization: EDD, WC, ACP. Funding acquisition: WC, ACP, EDD, MPK, ADA; Investigation: ES, EDD, JS, MS, WS, CD, AB, KWS, MPK, GP, LY, RJ, XL. Supervision: WC, ACP, EDD. Writing – original draft: ES, EDD; Writing – review & editing: ES, EDD, JS, ACP, WC, MPK, and KWS. Approving the final manuscript: All authors.

We would like to thank Dr Lesley Ellies for generously sharing the *Slc7a2^−/−^* mice with us. This research was supported by JDRF SRA-149-Q-R (to A.C.P. and E.D.D.), R01DK117147 (to W.C and A.C.P.), R01DK132669 (to E.D.D.) and K01DK117969 (to E.D.D.), by the University of Wisconsin–Madison, Department of Biochemistry and Office of the Vice Chancellor for Research and Graduate Education with funding from the Wisconsin Alumni Research Foundation (M.P.K.), and NIH grants, R01DK101573, R01DK102948, and RC2DK125961 (to A.D.A.). This research was performed using resources and/or funding provided by the NIDDK-supported Human Islet Research Network (UC4 DK104211 and DK112232) and by DK106755 (to A.C.P.), VUMC’s Digestive Diseases Research Center (P30 DK058404), and the Department of Veterans Affairs (BX000666). Experiments including confocal microscopy were performed in part using the Vanderbilt Cell Imaging Shared Resource (supported by NIH grants OD021630, CA68485, DK20593, DK58404, DK59637 and EY08126). Other technical assistance was provided by the Islet and Pancreas Analysis Core and the Hormone Assay and Analysis Core of Vanderbilt Diabetes Research and Training Center (P30 DK020593).

Dr. Danielle Dean is the guarantor of this work.

## Supplemental Figure Legends

**Supplemental Figure 1.**
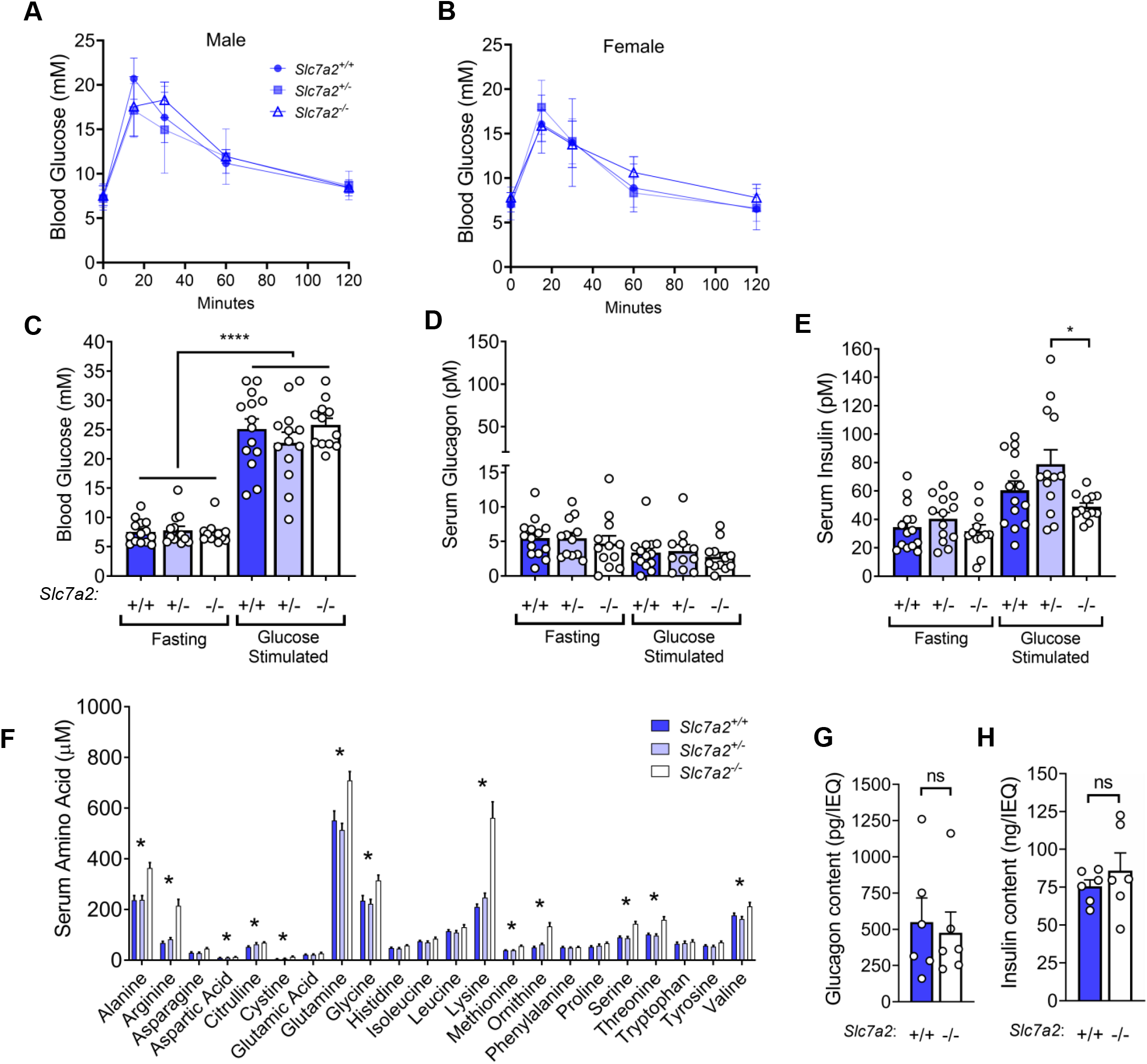
Characterization of *Slc7a2^−/−^* mice. (**A-B**) IP glucose tolerance tests in male and female *Slc7a2^+/+^* (n=3 each gender), *Slc7a2^+/−^* (n=6 each gender) and *Slc7a2^−/−^* (n=3 each gender) mice. (**C**) Blood glucose, **(D)** serum glucagon, and **(E)** serum insulin for *Slc7a2^+/+^*(n=14), *Slc7a2^+/−^* (n=12) and *Slc7a2^−/−^* (n=12) mice fasted for 6 hours, injected with *glucose* bolus and sampled 15 minutes post injection. (**F**) Analysis of serum amino acid concentrations in *Slc7a2^+/+^* (n=11), *Slc7a2^+/−^* (n=8) and *Slc7a2^−/−^* (n=12) mice. Asterisks indicate the 13 amino acids that were significantly higher in *Slc7a2^−/−^*animals than in *Slc7a2^+/+^* animals (p-value < 0.05). Total (**G**) glucagon and **(H)** insulin content of perifused islets (n=6 each) from **Fig 2 G, H.**

**Supplemental Figure 2.**
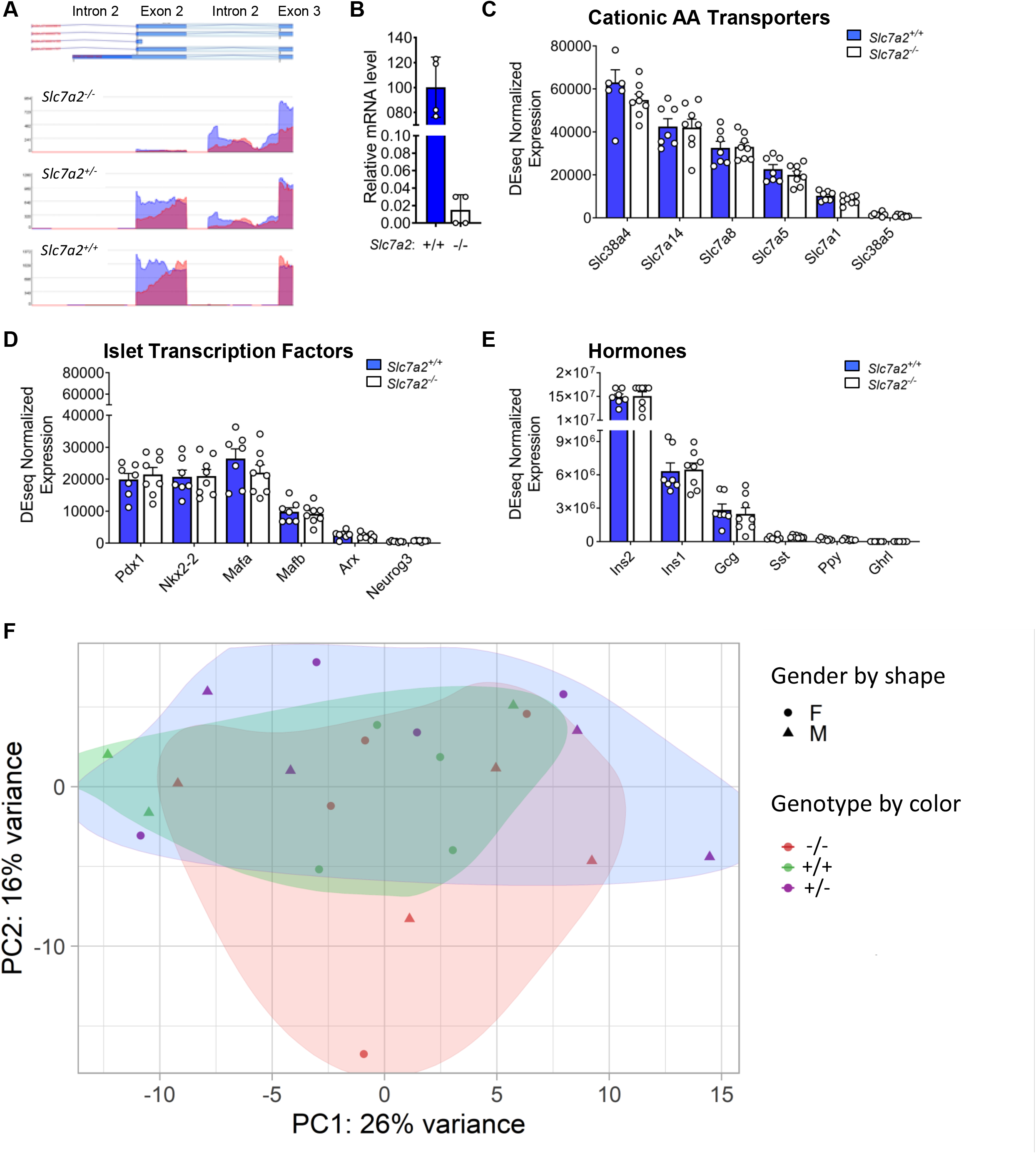
Loss of *Slc7a2* does not alter the islet gene expression profile. (**A**) Plot of individual RNAseq reads mapping to the *Slc7a2* locus from *Slc7a2^+/+^*, *Slc7a2^+/−^*and *Slc7a2^−/−^* islets. Stable mRNA product lacking Exon 2 sequence but containing sequence from Intron 2 was detectable in *Slc7a2^−/−^*islets. (**B**) Quantitative real-time RT-PCR analysis of *Slc7a2* Exon 2 in RNA isolated from *Slc7a2^−/−^* mice and *Slc7a2^+/+^* controls (n=4 each). (**C-E**) Comparison of islet-specific cationic amino acid transporter expression (**C**), islet transcription factor expression (**D**) and islet hormone expression (**E**) in *Slc7a2^+/+^* and *Slc7a2^-/^*^−^ mice by bulk RNAseq of isolated islets. (**F**) Principal component analysis of bulk RNAseq data from *Slc7a2^+/+^* and *Slc7a2^−/−^* mice showing clustering by gender, but not by genotype. (**G-H**) GO Term analyses of fold change data generated from gender clustered data.

**Supplemental Figure 3.**
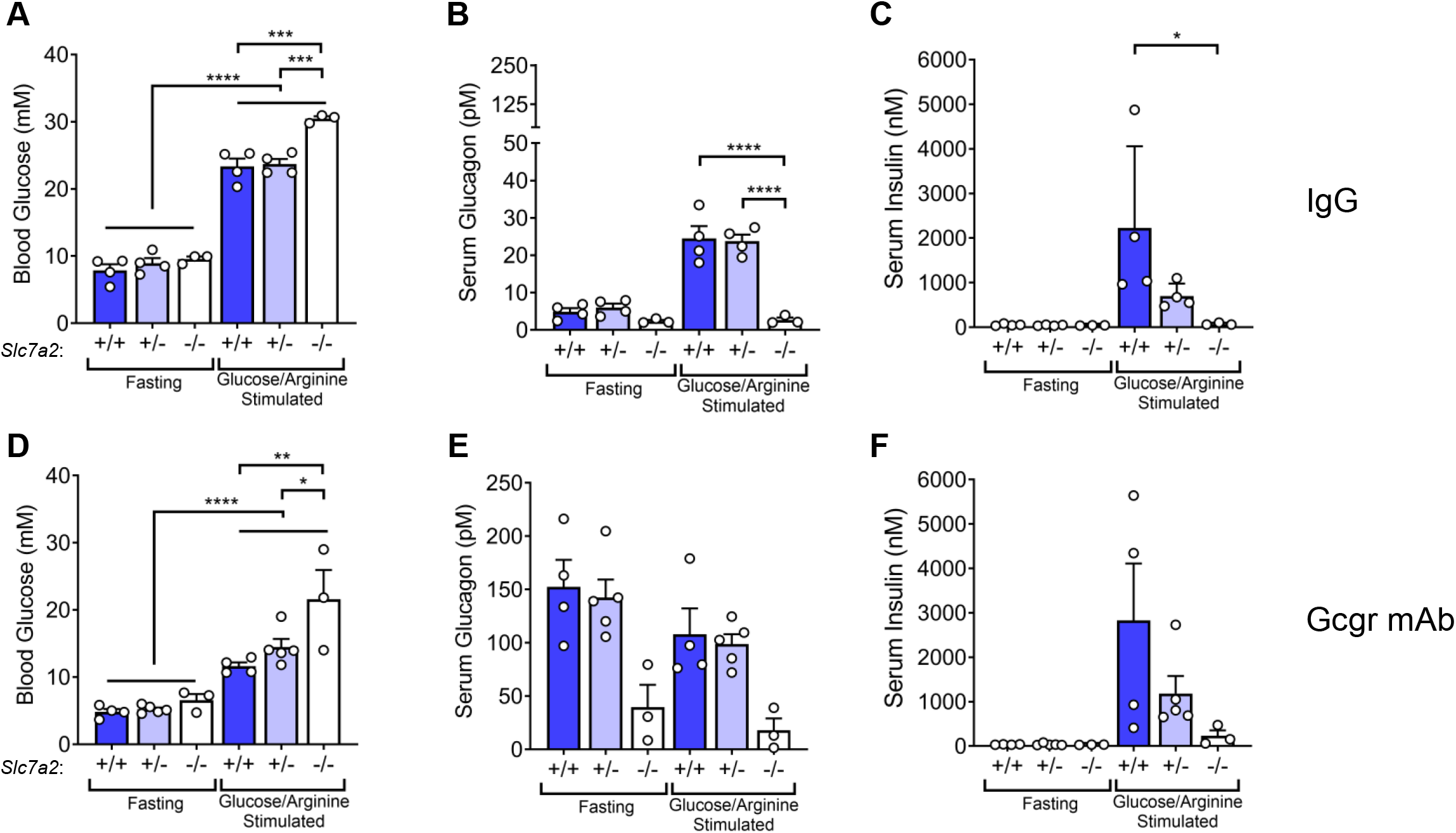
GCGR mAb-stimulated α cell proliferation does not alter stimulated glucagon or insulin secretion. (**A-C**) Blood glucose, serum glucagon and serum insulin for IgG treated *Slc7a2^+/+^* (n=4), *Slc7a2^+/−^*(n=4) and *Slc7a2^−/−^* (n=3) mice fasted for 6 hours, injected with glucose/arginine bolus and sampled 15 minutes post injection. (**D-F**) Blood glucose, serum glucagon and serum insulin for GCGR mAb treated *Slc7a2^+/+^*(n=4), *Slc7a2^+/−^* (n=5) and *Slc7a2^−/−^* (n=3) mice after glucose/arginine stimulation. All antibody treatments for two weeks prior to sampling serum.

## Supplemental Methods

### Intraperitoneal Glucose Tolerance Test

To assess the role of SLC7A2 in glucagon and glucose tolerance, *Slc7a2^+/+^*, *Slc7a2^+/−^* and *Slc7a2^−/−^* mice were fasted for 6 hours and then given intraperitoneal bolus of glucose to final concentrations of 2g / kg body weight. Blood was glucose was measured with a hand-held glucometer (Accu-Check Aviva) before injection (time 0) and at 15, 30, 60 and 120 minutes after injection.

### Serum amino acid analysis

Serum amino acids were analyzed in the Vanderbilt Hormone and Analytical Services core by HPLC. Serum samples were prepared by deproteinization with sulfosalicylic acid and adding lithium loading buffer to adjust the pH to an ideal level. Amino acids were separated using lithium-based ion exchange with ninhydrin post-column and detected on a Biochrom 30 amino acid analyzer. Peak results were analyzed to procure quantitative results.

### Whole Islet RNA sequencing

Whole Islets were pelleted and combined with to 20 μL lysis/binding solution in the RNAqueous micro-scale phenol-free total RNA isolation kit (Ambion). Trace DNA was removed with TURBO DNA-free (Ambion). RNA integrity was evaluated by Agilent 2100 Bioanalyzer. High integrity (RIN>7) total RNA was amplified, and cDNA libraries were constructed using the NEBNext® Poly(A) selection kit (New England Biolabs, Inc.). An Illumina NovaSeq 6000 was used to produce paired-end, 150-bp reads for each RNA sample yielding ∼1.7billion total raw reads. 8 replicates were included for groups *Slc7a2^+/−^* and *Slc7a2*^−/−^ whereas 7 replicates for *Slc7a2^+/+^*. Paired end raw reads were aligned to the reference mouse genome mm10 (GRCm38) using The Spliced Transcripts Alignment to a Reference (STAR) version 2.6 (1). ∼90% raw reads were uniquely mapped to genomic sites. Next, alignment quality and transcript quantification were performed using Strand NGS analysis platform 3.4 (Strand Life Sciences, India). Raw counts was normalized for library size and differential expression analysis using DESeq2 1.26 (2) with sex as covariate. Very lowly expressed genes were discarded from the analysis, keeping only genes covered by at least 20 reads in a minimum of three samples. Differentially expressed genes were defined by *P*_adj_ <0.05. RNA Sequencing data have been deposited in NCBI GEO with ID code GSE xxxxxx.. RNAseq analysis confirmed stable *Slc7a2* mRNA in *Slc7a2^−/−^* islets. Analysis of individual reads from RNAseq showed no reads in exon 2, the original knockout target, but some reads in intron 2 (Supplemental Fig 2A). Despite the presence of some RNA from this locus, the phenotypic results from these and previous studies indicate homozygous mutant mice are functionally null at the *Slc7a2* locus.

---

**Supplemental Table 1:** Amino acid concentrations in culture media for *ex vivo* islet α cell proliferation experiments.

| Amino Acids | RPMI 1640-based Media, Amino Acid Concentrations (μM) |  |  |  |  |  |  |  |  |  |
| --- | --- | --- | --- | --- | --- | --- | --- | --- | --- | --- |
|  | All AA - | All AA + | AIKMVOPSTYRHGN - | AIKMV - | OPSTY - | RHGN - | RHGN - / R + | RHGN - / H + | RHGN - / G + | RHGN - / N + |
| Glycine | 200 | 1500 | 200 | 1500 | 1500 | 200 | 200 | 200 | 1500 | 200 |
| L-Arginine | 40 | 600 | 40 | 600 | 600 | 40 | 600 | 40 | 40 | 40 |
| L-Asparagine | 40 | 350 | 40 | 350 | 350 | 40 | 40 | 40 | 40 | 350 |
| L-Aspartic acid | 10 | 10 | 10 | 10 | 10 | 10 | 10 | 10 | 10 | 10 |
| L-Cystine 2HCl | 10 | 10 | 10 | 10 | 10 | 10 | 10 | 10 | 10 | 10 |
| L-Glutamic Acid | 100 | 300 | 300 | 300 | 300 | 300 | 300 | 300 | 300 | 300 |
| L-Glutamine | 500 | 3250 | 3250 | 3250 | 3250 | 3250 | 3250 | 3250 | 3250 | 3250 |
| L-Histidine | 40 | 250 | 40 | 250 | 250 | 40 | 40 | 250 | 40 | 40 |
| L-Isoleucine | 125 | 250 | 125 | 125 | 250 | 250 | 250 | 250 | 250 | 250 |
| L-Leucine | 225 | 400 | 400 | 400 | 400 | 400 | 400 | 400 | 400 | 400 |
| L-Lysine hydrochloride | 200 | 1200 | 200 | 1200 | 1200 | 1200 | 1200 | 1200 | 1200 | 1200 |
| L-Methionine | 60 | 280 | 60 | 60 | 280 | 280 | 280 | 280 | 280 | 280 |
| L-Phenylalanine | 75 | 75 | 75 | 75 | 75 | 75 | 75 | 75 | 75 | 75 |
| L-Proline | 85 | 400 | 85 | 400 | 85 | 400 | 400 | 400 | 400 | 400 |
| L-Serine | 100 | 1250 | 100 | 1250 | 100 | 1250 | 1250 | 1250 | 1250 | 1250 |
| L-Threonine | 150 | 1750 | 150 | 1750 | 150 | 1750 | 1750 | 1750 | 1750 | 1750 |
| L-Tryptophan | 65 | 65 | 65 | 65 | 65 | 65 | 65 | 65 | 65 | 65 |
| L-Tyrosine | 50 | 250 | 50 | 250 | 50 | 250 | 250 | 250 | 250 | 250 |
| L-Valine | 300 | 600 | 300 | 300 | 600 | 600 | 600 | 600 | 600 | 600 |
| L-Alanine | 350 | 2250 | 350 | 350 | 2250 | 2250 | 2250 | 2250 | 2250 | 2250 |
| L-Ornithine | 100 | 400 | 100 | 400 | 100 | 400 | 400 | 400 | 400 | 400 |
| Data presented | Figures 1A, 4B and Supp. Figure 1A |  | Supp. Fig 1A |  |  | Figure 1A and Supp. Fig 1A | Figure 1A |  |  |  |

**Bold numbers** indicate high amino acid concentrations.

**Supplemental Table 2:**
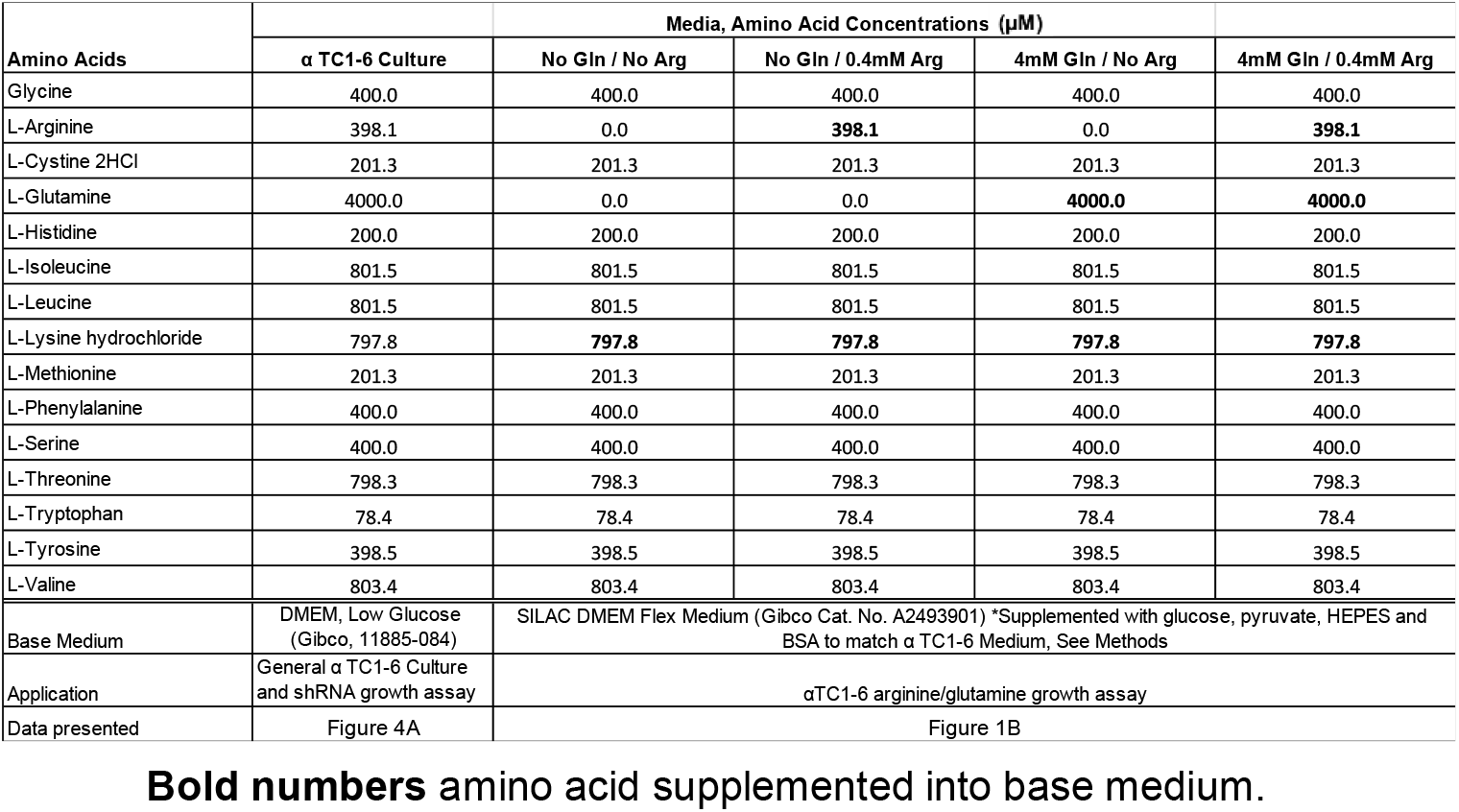
Amino acid concentrations in αTC1-6 culture media for cell growth assays.

| Amino Acids | Media, Amino Acid Concentrations (μM) |  |  |  |  |
| --- | --- | --- | --- | --- | --- |
|  | α TC1-6 Culture | No Gln / No Arg | No Gln / 0.4mM Arg | 4mM Gln / No Arg | 4mM Gln / 0.4mM Arg |
| Glycine | 400.0 | 400.0 | 400.0 | 400.0 | 400.0 |
| L-Arginine | 398.1 | 0.0 | **398.1** | 0.0 | **398.1** |
| L-Cystine 2HCl | 201.3 | 201.3 | 201.3 | 201.3 | 201.3 |
| L-Glutamine | 4000.0 | 0.0 | 0.0 | **4000.0** | **4000.0** |
| L-Histidine | 200.0 | 200.0 | 200.0 | 200.0 | 200.0 |
| L-Isoleucine | 801.5 | 801.5 | 801.5 | 801.5 | 801.5 |
| L-Leucine | 801.5 | 801.5 | 801.5 | 801.5 | 801.5 |
| L-Lysine hydrochloride | 797.8 | **797.8** | **797.8** | **797.8** | **797.8** |
| L-Methionine | 201.3 | 201.3 | 201.3 | 201.3 | 201.3 |
| L-Phenylalanine | 400.0 | 400.0 | 400.0 | 400.0 | 400.0 |
| L-Serine | 400.0 | 400.0 | 400.0 | 400.0 | 400.0 |
| L-Threonine | 798.3 | 798.3 | 798.3 | 798.3 | 798.3 |
| L-Tryptophan | 78.4 | 78.4 | 78.4 | 78.4 | 78.4 |
| L-Tyrosine | 398.5 | 398.5 | 398.5 | 398.5 | 398.5 |
| L-Valine | 803.4 | 803.4 | 803.4 | 803.4 | 803.4 |
| Base Medium | DMEM, Low Glucose (Gibco, 11885-084) | SILAC DMEM Flex Medium (Gibco Cat. No. A2493901) *Supplemented with glucose, pyruvate, HEPES and BSA to match α TC1-6 Medium, See Methods |  |  |  |
| Application | General α TC1-6 Culture and shRNA growth assay | αTC1-6 arginine/glutamine growth assay |  |  |  |
| Data presented | Figure 4A | Figure 1B |  |  |  |

**Bold numbers** amino acid supplemented into base medium.

**Supplemental Table 3:**
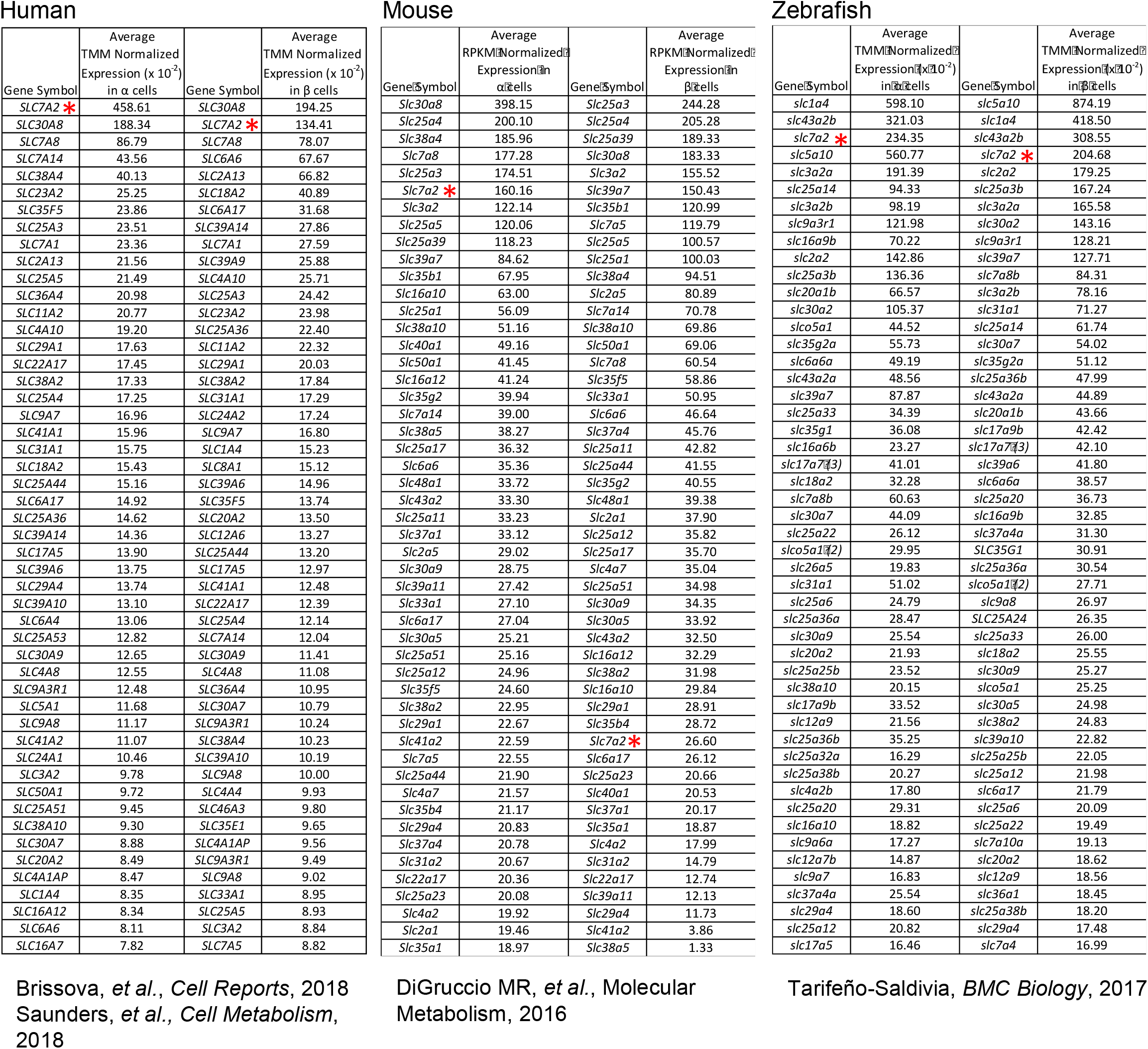
Top 50 solute carriers in Human, Mouse and Zebrafish α and β cells from highest to lowest relative expression.

## Notes

### Competing Interest Statement

All authors report no competing interests, except Kyle Sloop who is an employee of Lilly Inc.

## References

1. Wewer Albrechtsen NJ, Pedersen J, Galsgaard KD, Winther-Sorensen M, Suppli MP, Janah L, et al. The liver-alpha cell axis and type 2 diabetes. Endocr Rev. 2019.

2. Gelling RW, Du XQ, Dichmann DS, Romer J, Huang H, Cui L, et al. Lower blood glucose, hyperglucagonemia, and pancreatic alpha cell hyperplasia in glucagon receptor knockout mice. Proc Natl Acad Sci U S A. 2003;100(3):1438–43.

3. Johnson DG, Goebel CU, Hruby VJ, Bregman MD, and Trivedi D. Hyperglycemia of diabetic rats decreased by a glucagon receptor antagonist. Science. 1982;215(4536):1115-6.

4. Wang MY, Dean ED, Quittner-Strom E, Zhu Y, Chowdhury KH, Zhang Z, et al. Glucagon blockade restores functional beta-cell mass in type 1 diabetic mice and enhances function of human islets. Proc Natl Acad Sci U S A. 2021;118(9).

5. Kazda CM, Ding Y, Kelly RP, Garhyan P, Shi C, Lim CN, et al. Evaluation of Efficacy and Safety of the Glucagon Receptor Antagonist LY2409021 in Patients With Type 2 Diabetes: 12- and 24-Week Phase 2 Studies. Diabetes Care. 2016;39(7):1241–9.

6. Pettus J, Reeds D, Cavaiola TS, Boeder S, Levin M, Tobin G, et al. Effect of a glucagon receptor antibody (REMD-477) in type 1 diabetes: A randomized controlled trial. Diabetes Obes Metab. 2018;20(5):1302–5.

7. Kostic A, King TA, Yang F, Chan KC, Yancopoulos GD, Gromada J, et al. A first-in-human pharmacodynamic and pharmacokinetic study of a fully human anti-glucagon receptor monoclonal antibody in normal healthy volunteers. Diabetes Obes Metab. 2018;20(2):283–91.

8. Morgan ES, Tai LJ, Pham NC, Overman JK, Watts LM, Smith A, et al. Antisense Inhibition of Glucagon Receptor by IONIS-GCGRRx Improves Type 2 Diabetes Without Increase in Hepatic Glycogen Content in Patients With Type 2 Diabetes on Stable Metformin Therapy. Diabetes Care. 2019;42(4):585–93.

9. Kim J, Okamoto H, Huang Z, Anguiano G, Chen S, Liu Q, et al. Amino Acid Transporter Slc38a5 Controls Glucagon Receptor Inhibition-Induced Pancreatic alpha Cell Hyperplasia in Mice. Cell Metab. 2017;25(6):1348–61 e8.

10. Dean ED, Li M, Prasad N, Wisniewski SN, Von Deylen A, Spaeth J, et al. Interrupted Glucagon Signaling Reveals Hepatic alpha Cell Axis and Role for L-Glutamine in alpha Cell Proliferation. Cell Metab. 2017;25(6):1362–73 e5.

11. Solloway MJ, Madjidi A, Gu C, Eastham-Anderson J, Clarke HJ, Kljavin N, et al. Glucagon Couples Hepatic Amino Acid Catabolism to mTOR-Dependent Regulation of alpha-Cell Mass. Cell Rep. 2015;12(3):495–510.

12. Li M, Dean ED, Zhao L, Nicholson WE, Powers AC, and Chen W. Glucagon receptor inactivation leads to alpha-cell hyperplasia in zebrafish. J Endocrinol. 2015;227(2):93–103.

13. Wewer Albrechtsen NJ, Faerch K, Jensen TM, Witte DR, Pedersen J, Mahendran Y, et al. Evidence of a liver-alpha cell axis in humans: hepatic insulin resistance attenuates relationship between fasting plasma glucagon and glucagonotropic amino acids. Diabetologia. 2018;61(3):671–80.

14. Longuet C, Robledo AM, Dean ED, Dai C, Ali S, McGuinness I, et al. Liver-specific disruption of the murine glucagon receptor produces alpha-cell hyperplasia: evidence for a circulating alpha-cell growth factor. Diabetes. 2013;62(4):1196–205.

15. Akesson B, Henningsson R, Salehi A, and Lundquist I. Islet constitutive nitric oxide synthase and glucose regulation of insulin release in mice. J Endocrinol. 1999;163(1):39–48.

16. Henningsson R, and Lundquist I. Arginine-induced insulin release is decreased and glucagon increased in parallel with islet NO production. Am J Physiol. 1998;275(3):E500–6.

17. Fotiadis D, Kanai Y, and Palacin M. The SLC3 and SLC7 families of amino acid transporters. Mol Aspects Med. 2013;34(2-3):139–58.

18. Nicholson B, Manner CK, Kleeman J, and MacLeod CL. Sustained nitric oxide production in macrophages requires the arginine transporter CAT2. J Biol Chem. 2001;276(19):15881–5.

19. Manner CK, Nicholson B, and MacLeod CL. CAT2 arginine transporter deficiency significantly reduces iNOS-mediated NO production in astrocytes. J Neurochem. 2003;85(2):476–82.

20. Rothenberg ME, Doepker MP, Lewkowich IP, Chiaramonte MG, Stringer KF, Finkelman FD, et al. Cationic amino acid transporter 2 regulates inflammatory homeostasis in the lung. Proc Natl Acad Sci U S A. 2006;103(40):14895–900.

21. Singh K, Coburn LA, Barry DP, Asim M, Scull BP, Allaman MM, et al. Deletion of cationic amino acid transporter 2 exacerbates dextran sulfate sodium colitis and leads to an IL-17-predominant T cell response. Am J Physiol Gastrointest Liver Physiol. 2013;305(3):G225–40.

22. Brissova M, Haliyur R, Saunders D, Shrestha S, Dai C, Blodgett DM, et al. alpha Cell Function and Gene Expression Are Compromised in Type 1 Diabetes. Cell Rep. 2018;22(10):2667–76.

23. Saunders DC, Brissova M, Phillips N, Shrestha S, Walker JT, Aramandla R, et al. Ectonucleoside Triphosphate Diphosphohydrolase-3 Antibody Targets Adult Human Pancreatic beta Cells for In Vitro and In Vivo Analysis. Cell Metab. 2019;29(3):745–54 e4.

24. DiGruccio MR, Mawla AM, Donaldson CJ, Noguchi GM, Vaughan J, Cowing-Zitron C, et al. Comprehensive alpha, beta and delta cell transcriptomes reveal that ghrelin selectively activates delta cells and promotes somatostatin release from pancreatic islets. Mol Metab. 2016;5(7):449–58.

25. Tarifeno-Saldivia E, Lavergne A, Bernard A, Padamata K, Bergemann D, Voz ML, et al. Transcriptome analysis of pancreatic cells across distant species highlights novel important regulator genes. BMC Biol. 2017;15(1):21.

26. Singh K, Al-Greene NT, Verriere TG, Coburn LA, Asim M, Barry DP, et al. The L-Arginine Transporter Solute Carrier Family 7 Member 2 Mediates the Immunopathogenesis of Attaching and Effacing Bacteria. PLoS Pathog. 2016;12(10):e1005984.

27. Chen J, Spracklen CN, Marenne G, Varshney A, Corbin LJ, Luan J, et al. The trans-ancestral genomic architecture of glycemic traits. Nat Genet. 2021;53(6):840–60.

28. Pasquali L, Gaulton KJ, Rodriguez-Segui SA, Mularoni L, Miguel-Escalada I, Akerman I, et al. Pancreatic islet enhancer clusters enriched in type 2 diabetes risk-associated variants. Nat Genet. 2014;46(2):136–43.

29. Miguel-Escalada I, Bonas-Guarch S, Cebola I, Ponsa-Cobas J, Mendieta-Esteban J, Atla G, et al. Human pancreatic islet three-dimensional chromatin architecture provides insights into the genetics of type 2 diabetes. Nat Genet. 2019;51(7):1137–48.

30. Lee CS, Sund NJ, Behr R, Herrera PL, and Kaestner KH. Foxa2 is required for the differentiation of pancreatic alpha-cells. Dev Biol. 2005;278(2):484–95.

31. Conrad E, Dai C, Spaeth J, Guo M, Cyphert HA, Scoville D, et al. The MAFB transcription factor impacts islet alpha-cell function in rodents and represents a unique signature of primate islet beta-cells. Am J Physiol Endocrinol Metab. 2016;310(1):E91–E102.

32. Artner I, Le Lay J, Hang Y, Elghazi L, Schisler JC, Henderson E, et al. MafB: an activator of the glucagon gene expressed in developing islet alpha- and beta-cells. Diabetes. 2006;55(2):297–304.

33. Katoh MC, Jung Y, Ugboma CM, Shimbo M, Kuno A, Basha WA, et al. MafB Is Critical for Glucagon Production and Secretion in Mouse Pancreatic alpha Cells In Vivo. Mol Cell Biol. 2018;38(8).

34. Lawlor N, George J, Bolisetty M, Kursawe R, Sun L, Sivakamasundari V, et al. Single-cell transcriptomes identify human islet cell signatures and reveal cell-type-specific expression changes in type 2 diabetes. Genome Res. 2017;27(2):208–22.

35. Blachier F, Leclercq-Meyer V, Marchand J, Woussen-Colle MC, Mathias PC, Sener A, et al. Stimulus-secretion coupling of arginine-induced insulin release. Functional response of islets to L-arginine and L-ornithine. Biochim Biophys Acta. 1989;1013(2):144–51.

36. Dean ED. A Primary Role for alpha-Cells as Amino Acid Sensors. Diabetes. 2020;69(4):542–9.

37. Blachier F, Mourtada A, Sener A, and Malaisse WJ. Stimulus-secretion coupling of arginine-induced insulin release. Uptake of metabolized and nonmetabolized cationic amino acids by pancreatic islets. Endocrinology. 1989;124(1):134–41.

38. Fang Z, Weng C, Li H, Tao R, Mai W, Liu X, et al. Single-Cell Heterogeneity Analysis and CRISPR Screen Identify Key beta-Cell-Specific Disease Genes. Cell Rep. 2019;26(11):3132–44 e7.

39. Shrestha S, Saunders DC, Walker JT, Camunas-Soler J, Dai XQ, Haliyur R, et al. Combinatorial transcription factor profiles predict mature and functional human islet alpha and beta cells. JCI Insight. 2021;6(18).

40. Bosi E, Marchetti P, Rutter GA, and Eizirik DL. Human alpha cell transcriptomic signatures of types 1 and 2 diabetes highlight disease-specific dysfunction pathways. iScience. 2022;25(10):105056.

41. Yudkin JS, Forrest RD, Jackson CA, Ryle AJ, Davie S, and Gould BJ. Unexplained variability of glycated haemoglobin in non-diabetic subjects not related to glycaemia. Diabetologia. 1990;33(4):208–15.

42. Keller MP, Gatti DM, Schueler KL, Rabaglia ME, Stapleton DS, Simecek P, et al. Genetic Drivers of Pancreatic Islet Function. Genetics. 2018;209(1):335–56.

43. Mitok KA, Freiberger EC, Schueler KL, Rabaglia ME, Stapleton DS, Kwiecien NW, et al. Islet proteomics reveals genetic variation in dopamine production resulting in altered insulin secretion. J Biol Chem. 2018;293(16):5860–77.

44. Gerich JE, Charles MA, and Grodsky GM. Characterization of the effects of arginine and glucose on glucagon and insulin release from the perfused rat pancreas. J Clin Invest. 1974;54(4):833–41.

45. Capozzi ME, Svendsen B, Encisco SE, Lewandowski SL, Martin MD, Lin H, et al. beta Cell tone is defined by proglucagon peptides through cAMP signaling. JCI Insight. 2019;4(5).

46. Svendsen B, Larsen O, Gabe MBN, Christiansen CB, Rosenkilde MM, Drucker DJ, et al. Insulin Secretion Depends on Intra-islet Glucagon Signaling. Cell Rep. 2018;25(5):1127–34 e2.

47. Takahara T, Amemiya Y, Sugiyama R, Maki M, and Shibata H. Amino acid-dependent control of mTORC1 signaling: a variety of regulatory modes. J Biomed Sci. 2020;27(1):87.

48. Coburn LA, Singh K, Asim M, Barry DP, Allaman MM, Al-Greene NT, et al. Loss of solute carrier family 7 member 2 exacerbates inflammation-associated colon tumorigenesis. Oncogene. 2019;38(7):1067–79.

49. Sun T, Bi F, Liu Z, and Yang Q. SLC7A2 serves as a potential biomarker and therapeutic target for ovarian cancer. Aging (Albany NY). 2020;12(13):13281–96.

50. Xia S, Wu J, Zhou W, Zhang M, Zhao K, Liu J, et al. SLC7A2 deficiency promotes hepatocellular carcinoma progression by enhancing recruitment of myeloid-derived suppressors cells. Cell Death Dis. 2021;12(6):570.

51. Rodriguez-Ruiz ME, Buque A, Hensler M, Chen J, Bloy N, Petroni G, et al. Apoptotic caspases inhibit abscopal responses to radiation and identify a new prognostic biomarker for breast cancer patients. Oncoimmunology. 2019;8(11):e1655964.

52. Furuyama K, Chera S, van Gurp L, Oropeza D, Ghila L, Damond N, et al. Diabetes relief in mice by glucose-sensing insulin-secreting human alpha-cells. Nature. 2019;567(7746):43-8.

53. Jun LS, Millican RL, Hawkins ED, Konkol DL, Showalter AD, Christe ME, et al. Absence of glucagon and insulin action reveals a role for the GLP-1 receptor in endogenous glucose production. Diabetes. 2015;64(3):819–27.

54. Brissova M, Aamodt K, Brahmachary P, Prasad N, Hong JY, Dai C, et al. Islet microenvironment, modulated by vascular endothelial growth factor-A signaling, promotes beta cell regeneration. Cell Metab. 2014;19(3):498–511.

55. Kayton NS, Poffenberger G, Henske J, Dai C, Thompson C, Aramandla R, et al. Human islet preparations distributed for research exhibit a variety of insulin-secretory profiles. Am J Physiol Endocrinol Metab. 2015;308(7):E592–602.

## Supplemental References

1. Dobin A, Davis CA, Schlesinger F, Drenkow J, Zaleski C, Jha S, et al. STAR: ultrafast universal RNA-seq aligner. Bioinformatics. 2013;29(1):15–21.

2. Love MI, Huber W, and Anders S. Moderated estimation of fold change and dispersion for RNA-seq data with DESeq2. Genome Biol. 2014;15(12):550.

